# Neuronal lipid composition regulates distinct states of full-length tau assembly, mechanics and membrane interactions

**DOI:** 10.64898/2026.09.11.751000

**Authors:** Amaresh Kumar Mahakud, Aninda Sundar Modak, Snehasikta Singh, Vaishnavi Ananthanarayana, Hariharakrishnan Chidamabaram, Swagata Ghatak, Subashchandrabose Chinnathambi, Mohammed Saleem

## Abstract

Aggregation of tau into fibrillar assemblies and neurofibrillary tangles is a defining feature of tauopathies. However, the role of the membrane-rich neuronal environment on the assembly and mechanics of full-length tau remains unclear. Here, we examined how neuronal lipid membranes affect the assembly and membrane interaction of full-length human tau (hTau40). Lipid composition redirected hTau40 assembly, with total brain extract and brain phosphatidylcholine membrane producing shorter, more flexible fibrils, whereas brain phosphatidylserine membrane favoured a more heterogeneous and relatively rigid fibrillar population. Tau-membrane interaction depended strongly on the assembly state of tau. At the same protein concentration, preformed oligomers accumulated on membranes much more rapidly than monomeric tau, indicating that oligomer formation increases membrane recruitment. However, strong membrane binding did not directly predict the extent of membrane perturbation. Pronounced changes in lipid organisation were observed when tau was undergoing continued higher-order assembly at the membrane. Thus, the ability of tau to bind a membrane and its ability to reorganise that membrane appear to be related, but distinct, features of tau-membrane interaction. In cells, internalised hTau40 formed stable, low-mobility assemblies associated with the endolysosomal compartment and changes in its physical properties. These assemblies could also be transferred between cells in a membrane-derived vesicle model. Together, our results show that local membrane composition shapes tau assembly and mechanics, while the state of tau assembly influences how it binds to and remodels membranes.

## Introduction

The presence of neurofibrillary lesions and neuritic plaques in the cerebral cortex is the hallmark of Alzheimer’s disease (AD) [1, 2]. Despite the role of amyloid-β in the onset and progression of AD, the major pathology, including cognitive disruption, becomes more pronounced only after the tau inclusions start appearing. Dominantly inherited mutations are known to cause early-onset forms of neurodegenerative disease characterised by the accumulation of tau assemblies in the brain [2]. Although the molecular tau species responsible for neurodegeneration are unknown, short filaments are the main species of seed-competent tau in the brains of transgenic mice expressing human P301S tau [3]. The neurofibrillary lesions are comprised of paired helical filaments (PHFs) and straight tau filaments. The occurrence of tau fibrils in the diseased brain is explained by two mechanisms: first, nucleation of tau aggregation may occur inside the cell, driven by intracellular factors that remain largely elusive, without any input from neighbouring cells. Alternatively, tau assemblies propagate between cells, seeding aggregation in recipient cells in a “prion-like” manner. The observed spatiotemporal spread of tau misfolding in the human brain is consistent with observations of seeded aggregation in cultured cells[4–7] and in in vivo disease models, explained by this latter model [8–11]. Yet the cellular environments and their membrane interfaces that determine when soluble full-length tau (hTau40) enters an aggregation pathway, which assemblies are generated, and how those assemblies subsequently interact with cellular membranes remain poorly understood.

Monomeric tau, in its native state, is highly soluble and fails to self-assemble into fibrils under physiological conditions, even when hyperphosphorylated [12, 13]. Anionic lipid bilayers are potent inducers of tau nucleation and fibrillization[14], inducing helical structures and membrane disruption[15]. Tau oligomers are also known to impair phospholipid bilayer membrane integrity and cell viability[16]. The binding of the tau fragment K18 (a region within the microtubule binding repeat domain) to anionic membranes and micelles induces the formation of three distinct amphipathic helices located within repeat regions, which associate with the membrane surface[17–19]. Despite current understanding, a major limitation in previous studies has been the lack of complex neuronal membrane models. Another challenge has been capturing the comprehensive dynamics of tau nucleation, fibrillation, remodelling, and membrane deformation owing to the slow kinetics of tau aggregation.

Here, we investigate how neuronal membrane composition and tau assembly state jointly govern the reciprocal interaction between full-length human tau (hTau40) and lipid membranes. Using complementary reconstituted and cellular approaches, we examine whether neuronal lipid environments redirect tau aggregation and fibril mechanics, how monomeric, oligomeric, and actively assembling tau differ in membrane recruitment and remodelling, and whether these principles extend to intracellular membrane compartments following cellular uptake. Finally, we use a plasma-membrane-derived vesicle system to test whether persistent hTau40 assemblies remain compatible with membrane-mediated intercellular transfer. Together, this multiscale framework addresses how membrane composition, tau assembly state and membrane physical properties interact to shape the behaviour of full-length tau.

## Results

### Local neuronal lipid composition shapes distinct full-length tau assembly pathways

We first asked whether neuronal lipid composition alters the aggregation trajectory of full-length hTau40 (Fig. 1A). In the absence of heparin, hTau40 showed no detectable increase in ThT fluorescence during the ∼120h observation period, whereas the addition of heparin(Hep) produced a characteristic time-dependent increase in amyloid-associated fluorescence. Co-incubation with various brain-derived lipid extracts profoundly altered both the amplitude and kinetic trajectory of the aggregation curve, revealing a marked lipid-specific dependency (Table 1). Total brain lipid extract (TBE+hTau40+Hep) membrane reduced the endpoint ThT signal relative to the heparin-containing hTau40 control, lowering the plateau to ∼105 a.u. and increasing the rate of aggregation. In striking contrast, brain phosphatidylcholine (BPC+ hTau40+Hep) and a 6:4 ratio of brain phosphatidylcholine and brain phosphatidylserine (BPC-BPS+hTau40+Hep) membrane both accelerated the early aggregation kinetics, exhibiting steeper initial slopes and yielded higher final plateaus (∼130-135 a.u.), approaching the level of the heparin-only control. Most significantly, brain phosphatidylserine (BPS + hTau40 + Hep) produced the lowest ThT signal among the lipid conditions tested.

**Figure 1.**
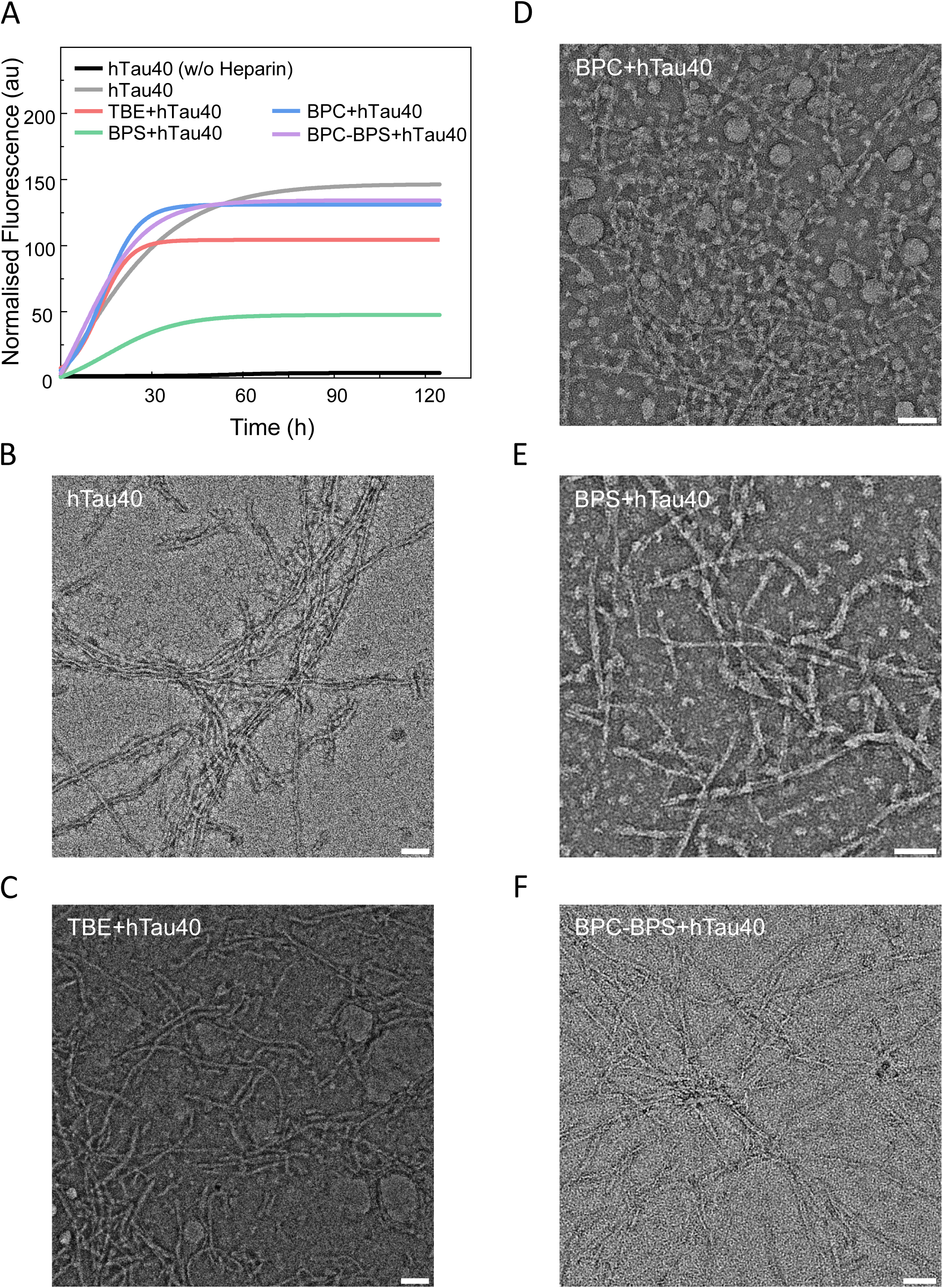
Neuronal membrane environment alters hTau40 aggregation kinetics and fibril morphology. **(A)** Thioflavin T (ThT) fluorescence kinetics of hTau40 (50 µM) aggregation in the presence of different lipid vesicle conditions, including TBE (Total brain extract), BPC (Brain phosphatidylcholine), BPS (Brain phosphatidylserine), and BPC: BPS (6:4), at a protein to lipid molar ratio of 1:4. Aggregation was monitored over 125 h. Reactions were induced with heparin (1:4 heparin to protein molar ratio), except for one control performed in the absence of heparin. Curves represent fitted normalised fluorescence data from three independent experiments (n = 3). **(B-F)** Representative transmission electron microscopy (TEM) micrographs of hTau40 aggregates at the endpoint of the aggregation reactions are shown in (A), with respective conditions labelled on the corresponding micrographs. Scale bar: 100 nm.

**Table 1.** hTau40 aggregation kinetics parameters. Growth rate and saturation fluorescence values derived from Boltzmann sigmoidal fits to normalised ThT fluorescence kinetics curves (Fig. 1A) for hTau40 aggregation alone and in the presence of TBE, BPC, BPS, and BPC-BPS membranes. Values represent fitted parameters ± standard error (SE).

| Membrane Conditions | Growth rate (au/h) | Saturation fluorescence (au) |
| --- | --- | --- |
| hTau40 | $3.8 \pm 0.7$ | $146.7 \pm 2.7$ |
| TBE+hTau40 | $5.2 \pm 0.8$ | $104.3 \pm 2.2$ |
| BPC+hTau40 | $5.9 \pm 0.6$ | $131 \pm 1.8$ |
| BPS+hTau40 | $1.2 \pm 0.6$ | $47.6 \pm 0.6$ |
| BPC-BPS+hTau40 | $4.7 \pm 0.5$ | $134.2 \pm 1.5$ |

As ThT fluorescence can be influenced by aggregate structure, dye accessibility, and aggregate abundance, we next examined endpoint morphology using negative-stain transmission electron microscopy (Fig. 1B-F). The hTau40+Hep condition displayed abundant, long, bundled filaments (Fig. 1B). In the presence of TBE lipid membrane, hTau40+Hep produced notably shorter and more dispersed fibrils (Fig. 1C). Intriguingly, despite exhibiting a substantial ThT signal, the BPC+hTau40+Hep samples were predominantly composed of small, globular particulate species and short fibrils (Fig. 1D). This finding suggests that the elevated ThT fluorescence observed under this condition may partly arise from non-fibrillar oligomeric or micellar assemblies rather than from canonical amyloid fibrils, in line with previous reports [16, 17]. The hTau40+Hep mixture in the presence of BPC-BPS membrane, however, closely recapitulated the extensive filamentous morphology of the heparin-only control (Fig. 1E). BPS+hTau40+Hep yielded the most heterogeneous pool of filaments (Fig. 1F). Collectively, these data hint that neuronal lipid composition redirects hTau40 into structurally distinct assembly states, influencing not only the extent of assembly but also the structural nature of the resulting species.

### Lipid composition determines the bending mechanics of hTau40 filaments

To determine whether membrane composition alters the mechanical properties of tau fibrils, we quantified the fibril contour length (L) and end-to-end distance (R) from TEM images and analysed them using a two-dimensional worm-like chain (2D WLC) model[20]. The resulting persistence length (*L*_*p*_) quantifies resistance to fibril bending, with larger *L*_*p*_values indicating greater bending stiffness (Fig. 2; Table 2)[21]. In the heparin-only hTau40 condition, fibrils exhibited a *L*_*p*_ of 218.1±65.8 nm (N=236), establishing the mechanical reference for the membrane-free condition (Fig. 2A). In the presence of TBE and BPC membranes, *L*_*p*_decreased markedly to 89.9±45.6 nm (N=670) and 94.9±41.2 nm (N=366), respectively (Fig. 2B, C). These values correspond to reductions of ∼57% and 56% relative to hTau40 alone, indicating substantially greater fibril flexibility in these membrane environments. The corresponding bending moduli, calculated from the persistence lengths, were 9.3±2.8, 3.8±1.9, and 4±1.7 (× 10^-28^) N·m² for hTau40, TBE+hTau40, and BPC+hTau40, respectively (Table 2), reflecting the same reduction in bending stiffness. In contrast, BPS-containing samples exhibited a higher

**Figure 2.**
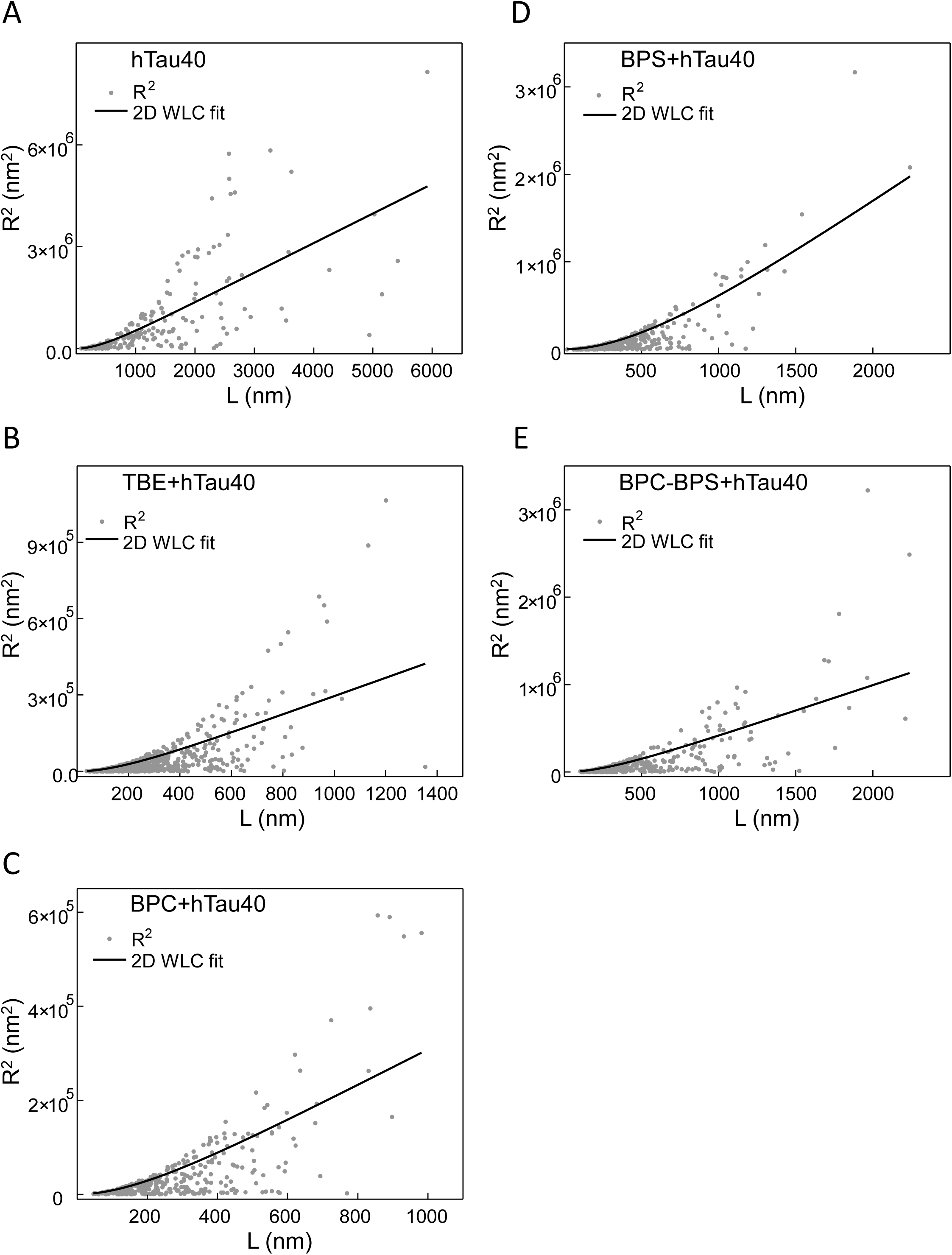
Neuronal membrane environment alters the flexibility of hTau40 fibrils. **(A-E)** Scatter plots of mean squared end-to-end distance (R²) versus contour length (L) for hTau40 fibrils formed under the indicated conditions, measured from TEM micrographs shown in Figure 1. Data were fitted using the 2D worm-like chain (WLC) model (solid line) to estimate the persistence length of fibrils. Respective conditions are labelled on individual panels. Persistence length (*L*_*p*_) values and number of fibrils analysed for each condition are summarised in Table 2.

**Table 2.**
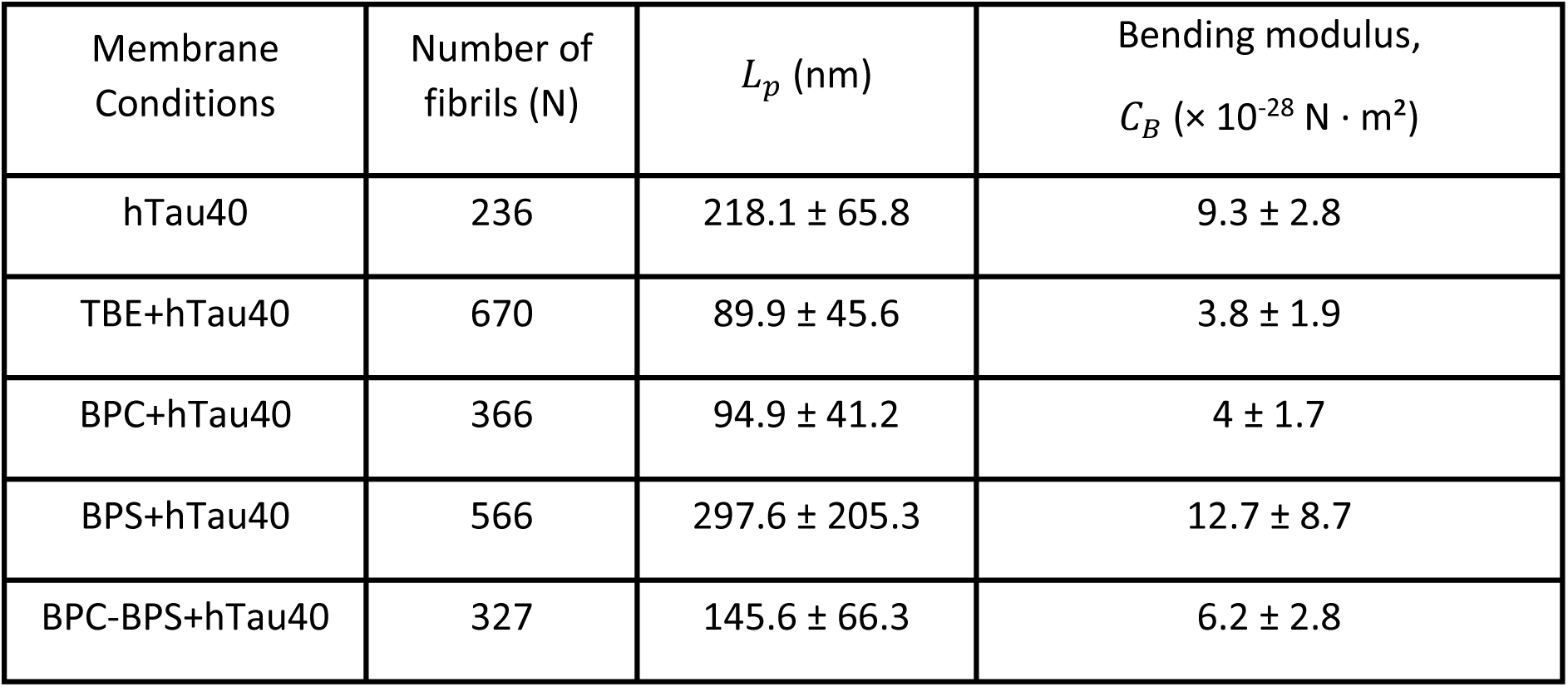
hTau40 fibril mechanical properties. Persistence length (*L*_*p*_) and bending modulus (*C*_*B*_) of hTau40 fibrils formed alone or in the presence of TBE, BPC, BPS, and BPC-BPS membranes, determined by two-dimensional worm-like chain (2D WLC) analysis of TEM images (Fig. 1B-F, 2). Uncertainty was estimated by bootstrap resampling (1000 iterations) and is reported as the mean ± half-width of the 95% confidence interval. The number of fibrils analysed for each condition is indicated in the table.

fitted persistence length of 297.6±205.3 nm (N=566; Fig. 2D). This corresponded to a bending modulus of 12.7±8.7 (× 10^-28^) N·m² (Table 2), the highest among the conditions examined. However, the large uncertainty associated with this estimate indicates considerable variation in fibril mechanical properties within this population. Thus, although the fitted whole population *L*_*p*_ was higher than that of the heparin-only condition, the large dispersion suggests that the BPS-associated fibrils cannot be adequately characterised by a uniform mechanical response. The BPC-BPS mixture (6:4) produced an intermediate persistence length of 145.6±66.3 nm (N=327; Fig. 2E), corresponding to a bending modulus of 6.2±2.8 (× 10^-28^) N·m² (Table 2), representing a ∼33% reduction relative to hTau40 alone, suggesting that the presence of anionic phosphatidylserine partially counteracts the effect of BPC alone. Together, these results demonstrate that membrane composition strongly influences the bending properties of tau fibrils, with TBE and BPC promoting more flexible fibrillar structures. In contrast, BPS produced a highly dispersed mechanical response.

To further resolve the mechanical diversity within each fibril population, we calculated the normalised bending parameter, η = (L−R)/L, for individual fibrils and represented it as a mass-weighted distribution of contour length (ΣL) across η bins (Fig. 3A-E). The experimental distributions showed substantial deviations from the theoretical 2D WLC distributions[20, 22] generated using the corresponding whole-population persistence lengths, indicating that a single persistence length does not fully capture the observed distributions of fibril bending behaviour. The hTau40 population displayed a broad distribution of η, with enrichment at both low η values (<0.2), corresponding to relatively straight fibrils, and higher η values (>0.4), corresponding to more strongly bent fibrils (Fig. 3A). TBE+hTau40 and BPC+hTau40 similarly exhibited broad distributions, with substantial contributions from intermediate η values (∼0.3-0.8; Fig. 3B, C). BPS+hTau40 showed the strongest deviation from its corresponding WLC prediction, particularly at η>0.2, despite its relatively high whole-population *L*_*p*_(Fig. 3D). The BPC-BPS+hTau40 condition displayed a broad distribution centred towards intermediate bending values, with a maximum around η≈0.4 (Fig. 3E). To determine whether these broad distributions reflected discrete fibril populations or broader mechanical heterogeneity, Gaussian mixture models containing one to six components were fitted to the individual-fibril η values, with model selection based on the Bayesian Information Criterion (BIC)[23]. The two-component model provided the lowest BIC for hTau40, with a ΔBIC of 37.2 relative to the single-component model, supporting the presence of two distinct mechanical populations. In contrast, the BIC favoured models with three to four components for TBE+hTau40, BPC+hTau40, BPS+hTau40, and BPC-BPS+hTau40, indicating substantially broader and more complex distributions of fibril-bending properties rather than a simple two-population organisation.

**Figure 3.**
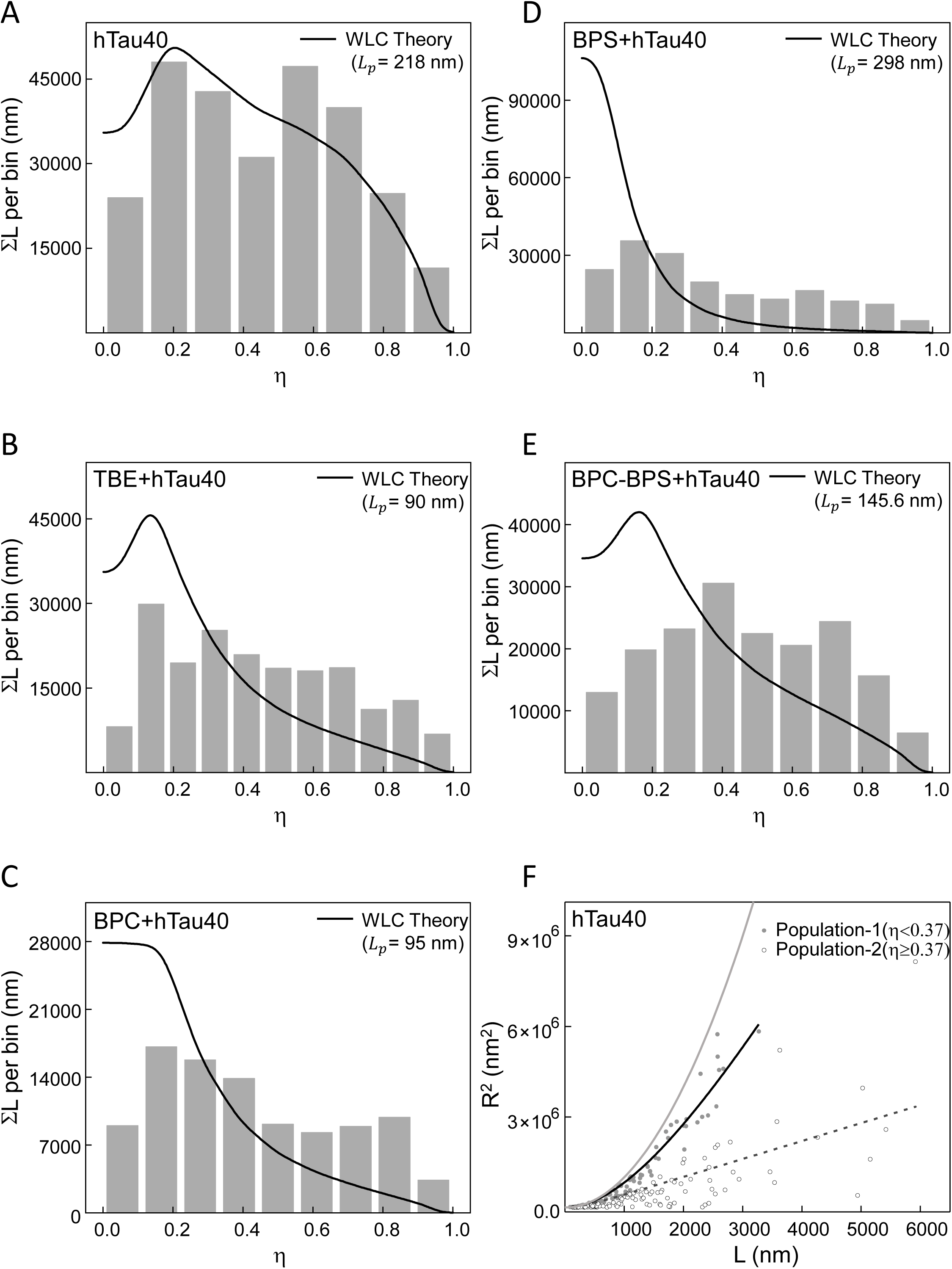
Neuronal membrane environment shifts hTau40 fibril bending flexibility distribution. **(A-E)** Histograms showing the mass-weighted distribution of summed contour length (ΣL per bin) as a function of the bending parameter η (η = (L − R)/L; see Results), where fibrils with η close to zero are relatively straight and rigid, and those with η close to one are highly curved and flexible. Solid lines represent the theoretical η distribution obtained from two-dimensional worm-like chain Monte Carlo simulations using the corresponding fitted persistence length (*L*_*p*_, indicated on individual panels). Respective conditions are labelled on individual panels. **(F)** Scatter plot of mean squared end-to-end distance (R²) versus contour length (L) for hTau40 fibrils (without membrane), segregated into two populations based on the bending parameter η: Population 1 (η < 0.37) and Population 2 (η ≥ 0.37), each fitted independently using the 2D WLC model (dashed and solid lines, respectively). The light grey line represents the theoretical limiting curve for ideally straight fibrils (R² = L²), which corresponds to the upper bound where the contour length and end-to-end distance coincide. The η cutoff corresponds to the minimum between the two peaks of the bimodal η distribution.

Within the membrane-free hTau40 population (Fig. 3A), the crossover between the two fitted Gaussian components occurred at η*=0.37, allowing the fibrils to be separated into a lower-η population (Population 1, N=121) and a higher-η population (Population 2, N=115). Independent 2D WLC analysis revealed a pronounced difference in persistence length between these populations. Population 1 exhibited a persistence length of 816±188 nm, whereas Population 2 exhibited a substantially lower persistence length of 149±63 nm (Fig. 3F), compared with the unsplit hTau40 population, with *L*_*p*_ = 218±66 nm. Consistent with these values, Population 1 fibrils clustered closer to the rigid-rod limit in the R^2^ versus L plot, whereas Population 2 contained fibrils with substantially smaller end-to-end distances relative to their contour lengths (Fig. 3F). Together, these findings indicate that neuronal membrane involvement fundamentally alters the mechanical organisation of tau fibrils. In the absence of membranes, hTau40 fibrils segregated into two populations with clearly distinct mechanical properties, whereas association with neuronal membranes produced broader and more complex distributions without a consistent discrete fibril population pattern. Thus, membrane association appears to disrupt the distinct mechanical partitioning observed in membrane-free tau fibrils, promoting a more heterogeneous mechanical landscape.

### Membrane composition and tau assembly state govern tau-membrane association and lipid-order perturbation

Having established that lipid composition influences tau aggregation behaviour, fibril population distribution, and their respective mechanical properties, we next examined how hTau40 engages a bilayer directly and in real time. We reconstituted giant unilamellar vesicles (GUVs) composed of complex neuronal lipids to mimic different local heterogeneities as described earlier. Treatment of hTau40-Alexa488 with the TBE membrane revealed progressive accumulation of tau on the membrane interface, first becoming evident at ∼8 h, prominently visible at ∼17 h and continuing through the ∼60 h observation period (Fig. 4A). Quantification of membrane-associated fluorescence across three successive temporal windows (0-25 h, 25-45 h, and 45-61 h) showed a significant increase between each interval (*p* < 0.0001 for all pairwise comparisons), demonstrating sustained and progressive enrichment of tau at the membrane interface (Fig. 4E). Interestingly, the growing tau pool containing a mixed population of aggregation states showed fibrillar growth perpendicular to the membrane plane starting ∼20 h onwards (Movie S1). In contrast, GUVs composed of BPC or the BPC-BPS (6:4) mixture showed no membrane-associated tau over the corresponding ∼60 h observation period (Fig. 4B, C, S3). Thus, these lipid compositions did not support appreciable tau recruitment under the chosen experimental conditions. GUVs composed exclusively of BPS exhibited a markedly different response. hTau40-488 accumulation at the vesicle periphery was detectable within minutes and was followed by pronounced membrane deformation and rupture, with vesicles losing their spherical morphology and undergoing disruption within ∼30 min (Fig. 4D, S2, Movie S2).

**Figure 4.**
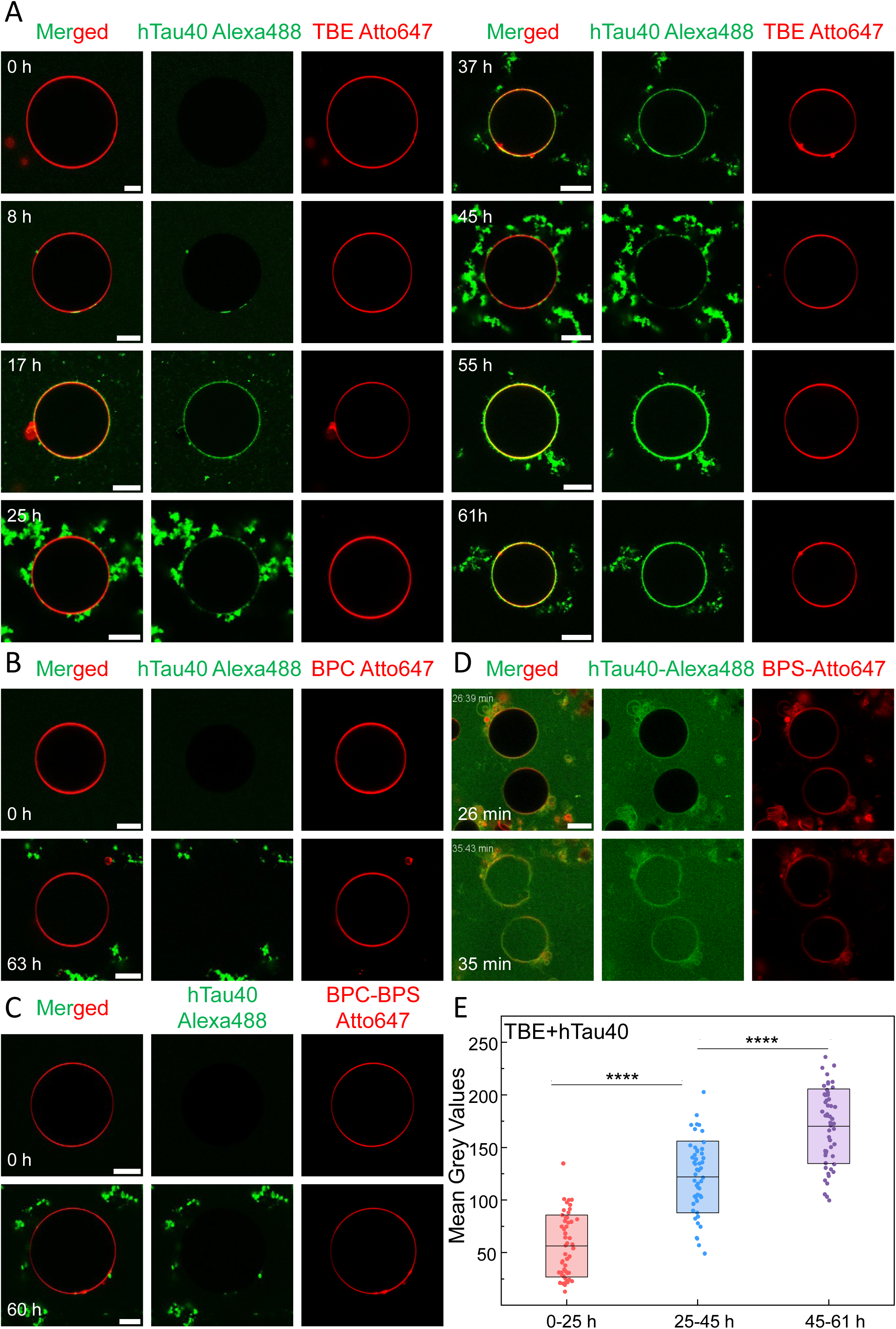
Progressive and selective membrane binding of hTau40 with neuronal lipid membranes. **(A)** Representative time-lapse confocal images showing the progressive binding of hTau40 (10 µM, Alexa Fluor 488, green) with TBE GUVs (Atto647, red) over a period of up to 3 days (∼ 60 h). Merged images together with the individual hTau40-Alexa488 and TBE-Atto647 fluorescence channels are shown at the indicated time points. **(B-C)** Representative confocal images of hTau40 (10 µM, Alexa Fluor 488, green) incubated with **(B)** BPC GUVs and **(C)** BPC-BPS GUVs (Atto647, red), acquired using the same imaging protocol as described in **(A)** over a period of up to 3 days. No detectable membrane binding of hTau40 with the membranes of either GUV composition was observed throughout the imaging period. **(D)** Confocal images of hTau40-Alexa488 with BPS GUVs (Atto 647, red), showing rapid accumulation of tau at the GUV periphery within minutes of exposure, followed by pronounced membrane deformation and loss of vesicle integrity. Images were acquired at the indicated time points. **(E)** Quantitative analysis of hTau40 binding to TBE GUVs, determined from mean grey values obtained by 360-point oval intensity profiling along the GUV membrane. Data were grouped into three time intervals: day 1 (0-25 h), day 2 (25-45 h), and day 3 (45-61 h). Approximately 50 GUVs were analysed for each time interval. Membrane-associated hTau40 fluorescence increased progressively over time. Data are presented as mean ± SD box plots with individual data points overlaid. Statistical significance was assessed using the Kruskal-Wallis test followed by Dunn’s post hoc analysis; **** *p* < 0.0001. Scale bar: 10 µm. All GUV experiments were performed independently in triplicate (*n* = 3).

These observations indicate that membrane composition strongly influences both the kinetics and consequences of tau-membrane association. The rapid response observed with BPS, together with the absence of detectable binding to BPC and BPC-BPS (6:4), suggests that a high degree of anionic lipid enrichment can strongly promote tau recruitment, whereas intermediate anionic content is insufficient under these conditions. Surface potential measurements (Fig. S1) broadly followed the expected differences in anionic lipid content; however, TBE and BPC-BPS exhibited similarly negative surface potentials despite markedly different tau-binding behaviour. Thus, net surface charge alone does not fully explain the observed differences in membrane recruitment. The contrasting behaviour of the different membrane compositions suggests that tau association is influenced by additional features of membrane organisation beyond overall surface charge, potentially including lipid packing and headgroup presentation. Notably, the native-like complexity of TBE supported gradual and sustained tau association without the acute membrane disruption observed for BPS.

We next examined whether membrane association differed among distinct assembly states of hTau40 using TBE-containing GUVs. Pre-formed oligomeric Alexa647 hTau40 displayed progressive accumulation at the GUV membrane, with membrane-associated fluorescence detectable as early as ∼2 h and increasing substantially over the subsequent time points before approaching a plateau between ∼12 and ∼26 h (Fig. 5A), likely indicating the tau oligomers saturating the membrane interface in two-dimensions. Quantification of membrane-associated fluorescence across successive time windows confirmed a significant increase between the initial 0-12 h interval and the later intervals (*p* < 0.01; Fig. 6A, S5), demonstrating efficient and progressive recruitment of oligomeric tau to the membrane surface. In contrast, monomeric Alexa647 hTau40 showed no detectable membrane association throughout the 72 h observation period (Fig. 5B). The mature aggregate preparation exhibited a markedly different appearance, indicating binding at early time points evident from fluorescence intensity of the equatorial plane; however, we did not observe a clear increase in membrane-associated fluorescence or the progressive attachment of discrete aggregate structures to the vesicle surface (Fig. 5C). The early fluorescence observed in the aggregate preparation may therefore reflect membrane-active species present within the heterogeneous aggregate sample, potentially including oligomeric species.

**Figure 5.**
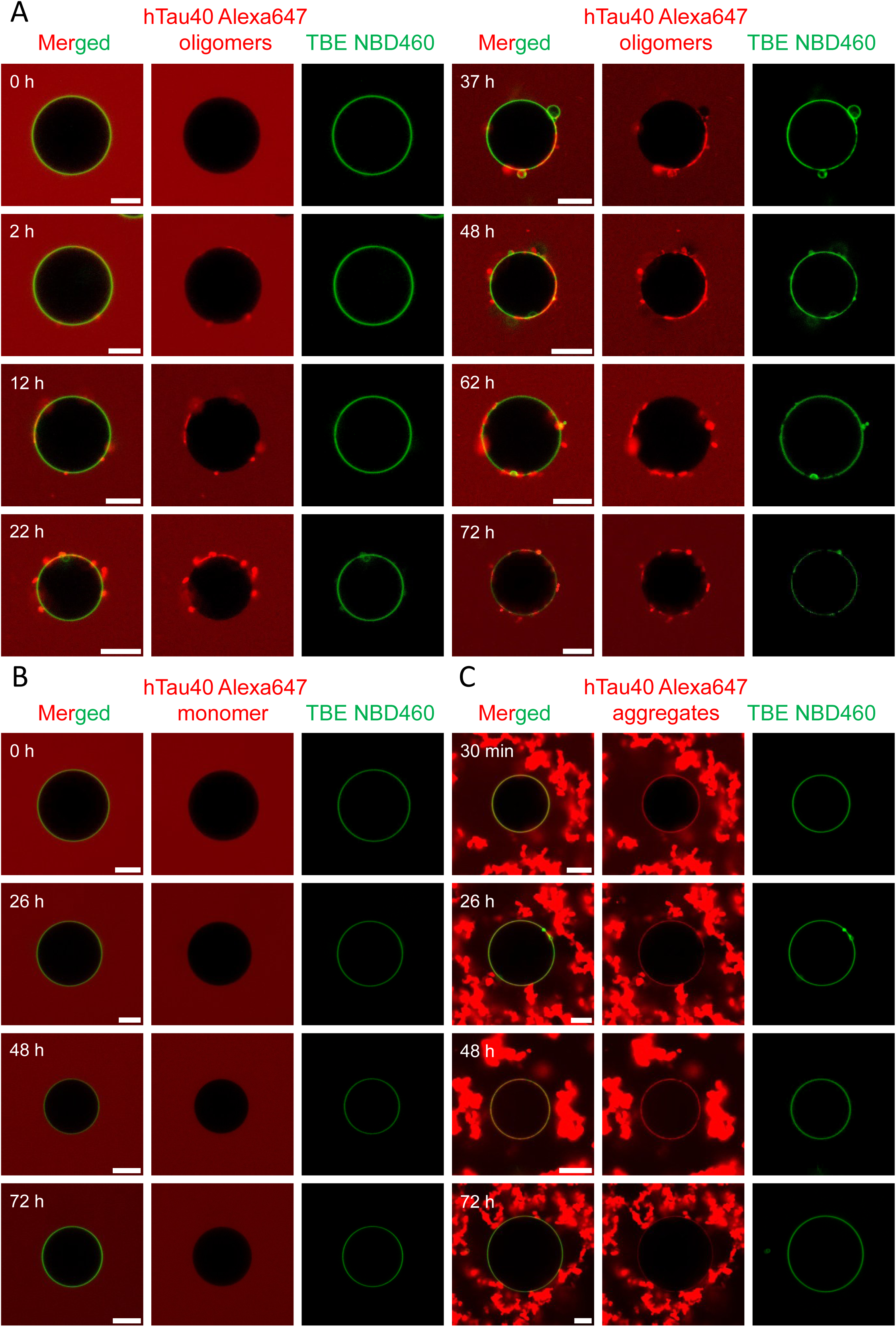
Selective membrane binding by hTau40 oligomers identifies them as the primary membrane-binding species. **(A)** Representative time-lapse confocal images showing the progressive binding of hTau40 oligomers(10µM, Alexa647, red) to TBE GUVs (NBD460, green) over a period of 72 h. Merged images together with the hTau40-Alexa647 channel are shown at the indicated time points. **(B)** Time-lapse confocal images of hTau40 monomers (10µM, Alexa647, red) incubated with TBE GUVs (NBD460, green) over the same 72 h period. No detectable membrane binding of hTau40 monomers was observed at any time point. **(C)** Time-lapse confocal images of hTau40 aggregates (10µM, Alexa647, red) incubated with TBE GUVs (NBD460, green) over the same 72 h period. In contrast to oligomers, aggregated hTau40 was detected at the membrane from the earliest time point (0 h), with no further increase in membrane-associated fluorescence during the course of the experiment. Membrane binding at the earliest time point may arise from residual oligomeric species in the aggregate preparation that retain membrane-binding capability; however, this possibility remains speculative. Scale bar: 10 µm. All GUV experiments were performed independently in triplicate (*n* = 3).

**Figure 6.**
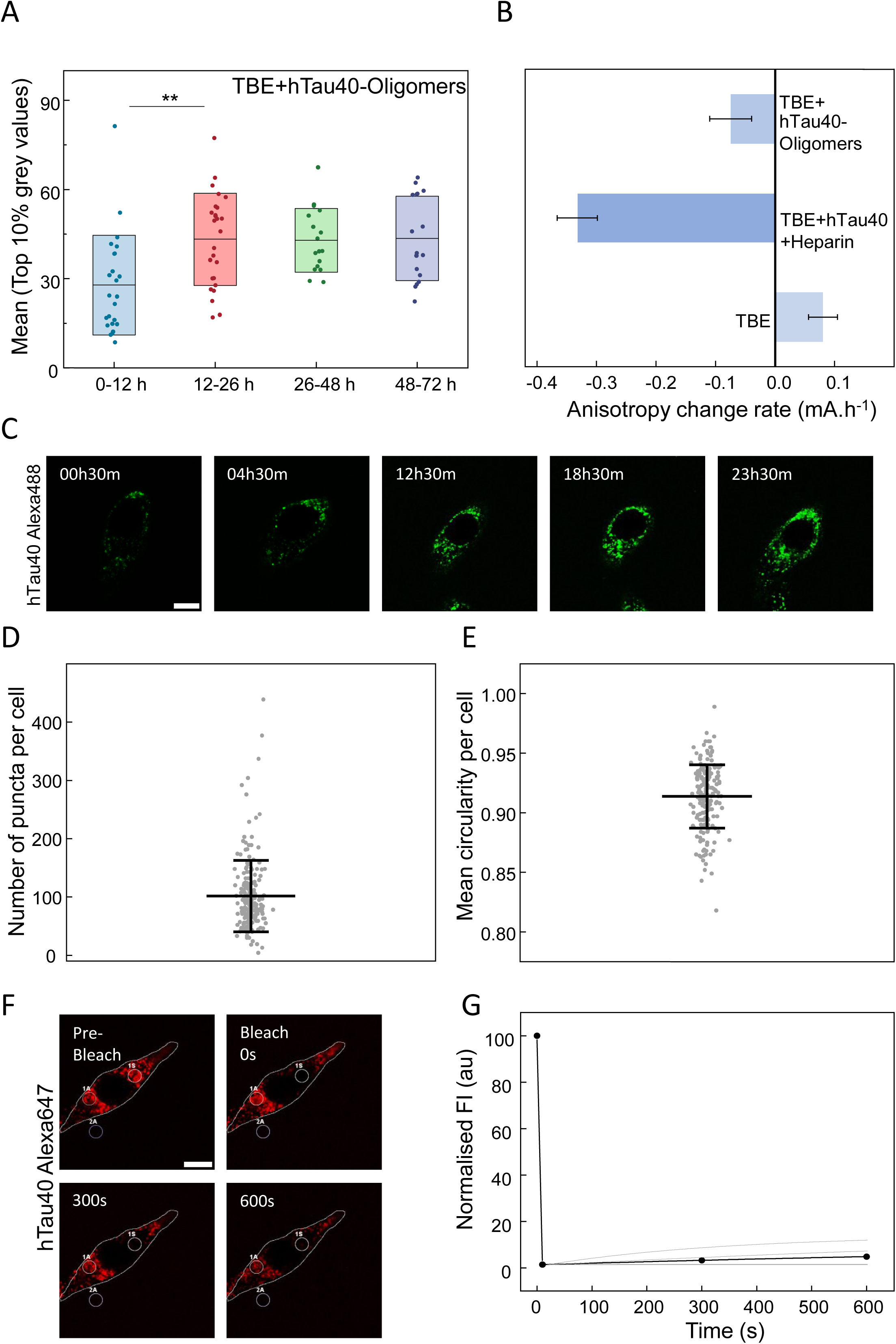
Oligomeric hTau40 binds to and perturbs neuronal lipid membranes, and extracellular hTau40 forms stable intracellular puncta following cellular uptake. (**A**) Quantification of hTau40 oligomer binding to TBE GUVs, determined from the mean of the top 10% grey values obtained by 360-point oval intensity profiling along the GUV membrane. Data were grouped into four time intervals (0-12 h, 12-26 h, 26-48 h and 48-72 h), with approximately 20 GUVs analysed per interval. Membrane-associated fluorescence increased significantly between 0-12 h and 12-26 h, consistent with progressive membrane accumulation during the early stages of incubation. Data are presented as mean ± SD box plots with individual data points overlaid. Statistical significance was determined by the Kruskal-Wallis test with Dunn’s post hoc analysis; ** *p* < 0.01. **(B)** Changes in membrane order were assessed by steady-state TMA-DPH fluorescence anisotropy using TBE LUVs incubated with full-length hTau40 (50µM with heparin; tau: heparin, 4:1) or hTau40 oligomers (1µM without heparin) (200 µM lipid; lipid: TMA-DPH ratio 100:1) over 125 h. The rate of anisotropy change was determined from the slope of a linear fit to anisotropy values over time, where positive and negative slopes indicate increasing and decreasing membrane order, respectively. Data are presented as mean ± SD from three independent experiments. **(C)** Representative time-lapse confocal images showing intracellular puncta formation in HT22 hippocampal cells following extracellular addition of hTau40-Alexa488 (1 µM) in the presence of heparin (tau: heparin, 4:1). Intracellular hTau40 puncta were detectable within 30 min of extracellular hTau40 addition, indicating early cellular uptake followed by progressive puncta formation over the 24 h imaging period. **(D-E)** Quantification of intracellular hTau40 puncta (D) number per cell and (E) circularity 24 h after extracellular hTau40 addition. Cells contained approximately 100 puncta per cell, with a mean circularity of ∼0.9, indicating a predominantly spherical puncta morphology. Data are presented as mean ± SD, pooled from three independent experiments (∼200 cells analysed in total). **(F-G)** Fluorescence recovery after photobleaching (FRAP) analysis of hTau40-Alexa647 puncta in HT22 cells, 24 h after extracellular hTau40 addition. **(F)** Representative confocal images before bleaching (pre-bleach), immediately after bleaching (0 s), and during recovery (300 s, 600 s). ROIs 1S, 1A, and 2A denote the bleached, reference, and background region, respectively. **(G)** Normalised fluorescence intensity recovery curves over time; grey traces represent recovery curves from five independent cell/region measurements, and the black trace represents the mean, showing negligible fluorescence recovery, indicative of mature, immobile puncta. Scale bar: 10 µm.

These distinct membrane-binding behaviours were accompanied by differential effects on membrane lipid order, as assessed by time-resolved fluorescence anisotropy measurements (Fig. 6B, S6). TBE membranes alone exhibited only a modest positive anisotropy drift over time (+0.08 mA h⁻¹), indicating comparatively stable membrane organisation over the experimental period. In contrast, TBE membranes incubated with initially monomeric hTau40 under heparin-induced aggregation conditions showed a pronounced decrease in anisotropy, with slopes of ∼ −0.32 to −0.35 mA h⁻¹, indicating substantial loss of lipid packing/order during ongoing tau aggregation. Pre-formed hTau40 oligomers produced a considerably smaller decrease in anisotropy (∼−0.08 mA h⁻¹), despite their substantially stronger membrane association. Importantly, the two tau conditions were selected to address distinct aspects of membrane remodelling. hTau40 was used at 50 µM to generate a reproducible heparin-induced aggregation reaction within an experimentally accessible timescale, allowing membrane-order changes to be examined during ongoing tau assembly. In contrast, pre-formed oligomers were tested at 1 µM to determine whether an assembly state with pronounced membrane affinity could perturb lipid organisation at substantially lower bulk protein exposure. Both conditions reduced membrane anisotropy, demonstrating that tau assembly, as well as preformed oligomers, can alter membrane organisation. Notably, oligomeric tau retained a measurable effect despite being present at a concentration 50-fold lower. Because the two conditions differ in both concentration and assembly state, their absolute slopes are interpreted as responses under distinct experimental regimes rather than as a direct comparison of intrinsic membrane-disordering potency. Collectively, these observations demonstrate that both ongoing hTau40 assembly and pre-formed oligomeric tau perturb neuronal membrane lipid organisation, albeit under distinct concentration regimes. This provides a mechanistic basis for investigating how tau-membrane interactions translate into cellular uptake, intracellular accumulation, and membrane-associated phenotypes.

### Internalised hTau40 forms stable, low-mobility puncta associated with changes in lysosomal membrane physical properties

The in vitro data above indicate that membrane perturbation is more strongly associated with ongoing tau fibrillization than with membrane binding alone. We next asked whether a similar distinction between dynamic and stable tau assemblies could be observed in living neuronal cells. Although liquid-liquid phase separation is increasingly recognised as an early and potentially reversible mode of tau organisation, the transition from dynamic condensates to stable, less mobile assemblies remains poorly understood. To examine whether a polyanionic cofactor could promote this transition, ht22 hippocampal neuronal cells were treated with heparin and hTau40-Alexa488. Intracellular tau puncta became detectable within 30 min and were readily observed after 24 h of treatment (Fig. 6C). Quantification at 24 h revealed ∼100 hTau40 puncta per cell, with a mean circularity of ∼0.9, indicating that the puncta were predominantly spherical (Fig. 6D, E). To determine the dynamic properties of these intracellular assemblies, fluorescence recovery after photobleaching (FRAP) of hTau40-Alexa647 was performed over 600 s. Following near-complete photobleaching, with normalised fluorescence decreasing from 100% to <5%, no measurable fluorescence recovery was observed throughout the acquisition period (Fig. 6F, G), indicating that the puncta were highly immobile on this timescale. This behaviour contrasts with tau assemblies previously reported in HT22 cells in the absence of heparin, which showed a significantly lower abundance of puncta exhibiting liquid-like characteristics and rapid fluorescence recovery [24]. Together, these observations indicate that exposure to a polyanionic cofactor such as heparin promotes the formation of abundant, predominantly spherical intracellular tau assemblies with markedly restricted molecular mobility, consistent with a transition toward a more stable, solid-like state rather than simply enhanced formation of dynamic condensates.

We next investigated the subcellular localisation and organellar interactions of these pathological aggregates. Strikingly, tau puncta exhibited substantial spatial overlap with LysoTracker-positive lysosomes (Fig. 7A), with Pearson’s correlation coefficient approaching 0.8 across cells (Fig. 7B), indicating a strong association of the intracellular tau assemblies with the lysosomal compartment. This association raised a critical mechanistic question: does the presence of solid-like tau aggregates merely reflect passive entrapment within lysosomes, or do they actively perturb the biophysical state of the lysosomal membrane? To distinguish between these possibilities, we employed fluorescence lifetime imaging microscopy (FLIM) in conjunction with two distinct environmentally sensitive probes targeted to lysosomes: a membrane-tension sensor (LysoFlipper^TR^) that senses lipid packing/order and a viscosity-sensitive probe (JIND-Mor)[25, 26]. These probes enable quantitative, live-cell readouts of organelle physical properties at nanoscale resolution. In untreated cells, LysoFlipper^TR^ exhibited a mean fluorescence lifetime of 4.9 ns, whereas hTau40-treated cells showed a significant reduction to 4.3 ns (*p* < 0.0001; Fig. 7C, D). Similarly, the JIND-Mor lifetime decreased from 2.8 ns in untreated cells to 2.6 ns following hTau40 treatment (*p* < 0.001; Fig. 7E, F). Thus, tau-containing puncta were not only associated with lysosomes but were accompanied by measurable changes in both lysosomal membrane lipid packing and viscosity.

**Figure 7.**
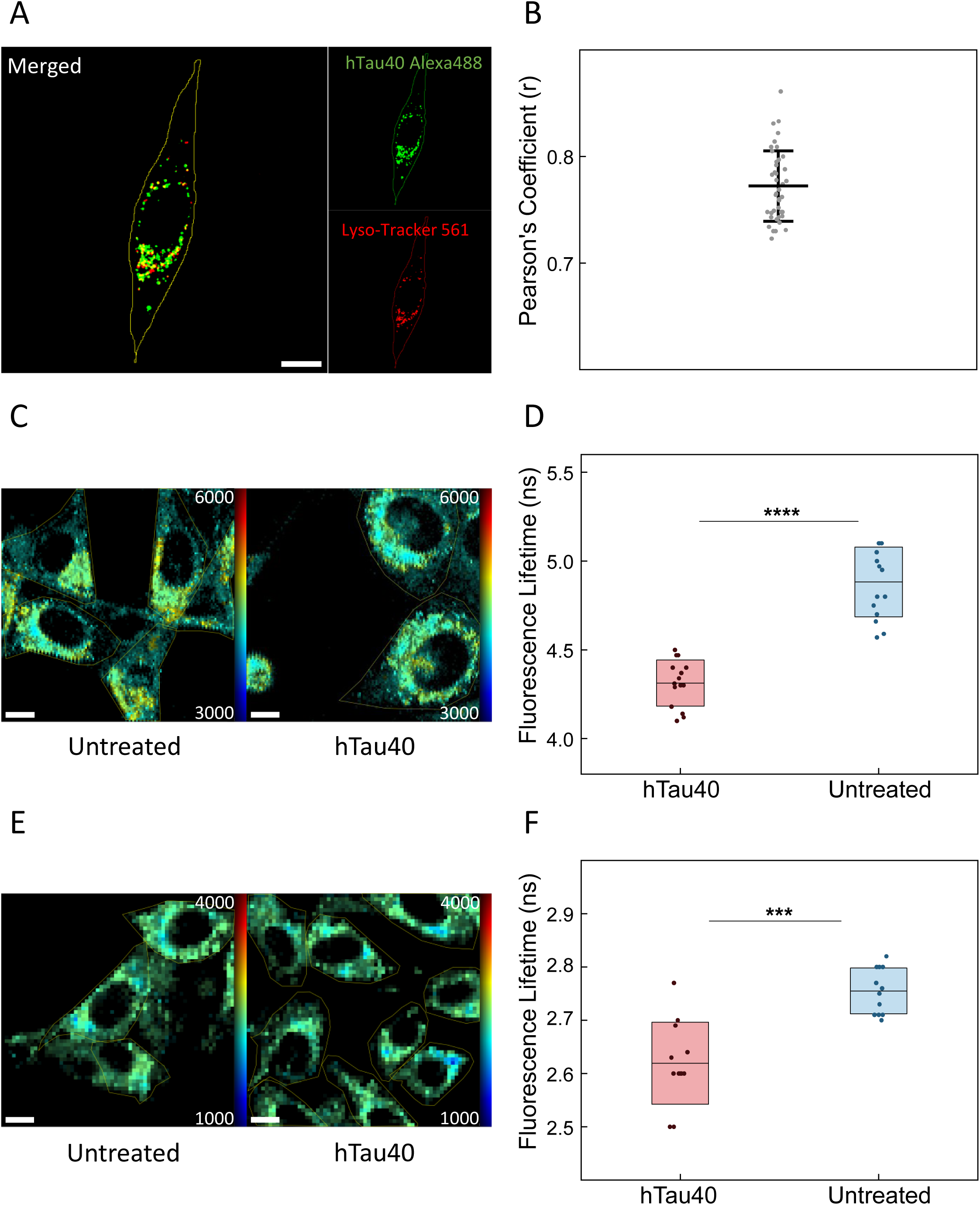
hTau40 colocalises with lysosomes and is associated with reduced lysosomal membrane order and viscosity. **(A-B)** HT22 cells were incubated with hTau40-Alexa488 (1 µM, tau: heparin ratio 4:1) and, following puncta formation, lysosomal colocalisation was assessed using LysoTracker 561. **(A)** Representative confocal images showing merged, hTau40-Alexa488, and LysoTracker channels. **(B)** Quantification of colocalisation using Pearson’s coefficient (r), demonstrating strong colocalisation (r ≈ 0.8) between hTau40 puncta and lysosomes. Data are presented as scatter dot plots with mean ± SD, from three independent experiments (∼40 regions analysed). **(C-D)** Impact of hTau40 on lysosomal membrane order, assessed by fluorescence lifetime imaging microscopy (FLIM) using Lyso Flipper-TR. **(C)** Representative FLIM images of untreated and hTau40-treated HT22 cells; colour scale represents fluorescence lifetime from 3000 to 6000 ps. **(D)** Quantification showed a significant reduction in fluorescence lifetime following hTau40 treatment compared with untreated controls, consistent with reduced lysosomal membrane order. Data are presented as mean ± SD box plots with individual data points overlaid, from three independent experiments; **** *p* < 0.0001. **(E-F)** Impact of hTau40 on lysosomal membrane viscosity, assessed by FLIM using the JIND-Mor viscosity-sensitive dye. **(E)** Representative FLIM images of untreated and hTau40-treated HT22 cells; colour scale represents fluorescence lifetime from 1000 to 4000 ps. **(F)** Quantification showed a significant reduction in fluorescence lifetime in hTau40-treated cells compared with untreated controls, consistent with reduced lysosomal membrane viscosity. Data are presented as mean ± SD box plots with individual data points overlaid, from three independent experiments; *** *p* < 0.001. Statistical significance was determined using the Kruskal-Wallis test with Dunn’s post hoc analysis. Scale bar: 10 µm.

### Membrane-derived vesicles enable intercellular transfer of hTau40 assemblies

Having established that extracellularly supplied hTau40 forms stable, low-mobility intracellular puncta associated with the endolysosomal compartment, we next asked whether these assemblies could be captured within a plasma-membrane-derived vesicular compartment and transferred to recipient cells. We employed giant plasma membrane vesicles (GPMVs), experimentally induced plasma membrane vesicles that retain much of the lipid and protein complexity of the parental plasma membrane and can enclose cytoplasmic material during vesiculation (Fig. 8A, schematic). GPMVs were generated from HT22 donor cells containing hTau40-Alexa488 puncta. Following isolation, a subset of GPMVs retained punctate hTau40 fluorescence (Fig. 8B, C, Movie S3, S4), indicating association of intracellular tau assemblies with the GPMV fraction and consistent with their capture during plasma-membrane vesiculation. Previous studies have shown that GPMVs can incorporate cytoplasmic material and even preloaded particulate cargo during their formation, providing a physical basis for the entrapment of intracellular assemblies[27]. Building on this observation, we next asked whether GPMV-packaged tau could functionally deliver its cargo into recipient cells (Fig. 8A, schematic). When hTau40-containing GPMVs were subsequently applied to untreated recipient HT22 cells, intracellular hTau40-Alexa488 puncta were detected after 24 h (Fig. 8D). Thus, membrane-associated hTau40 present in the GPMV preparation remained competent for transfer to recipient cells. The persistence of punctate rather than diffuse fluorescence following transfer suggests that the delivered protein remained associated with a concentrated assembly state. However, the present experiments do not distinguish between the persistence of the transferred donor assembly and reorganisation following cellular uptake.

**Figure 8.**
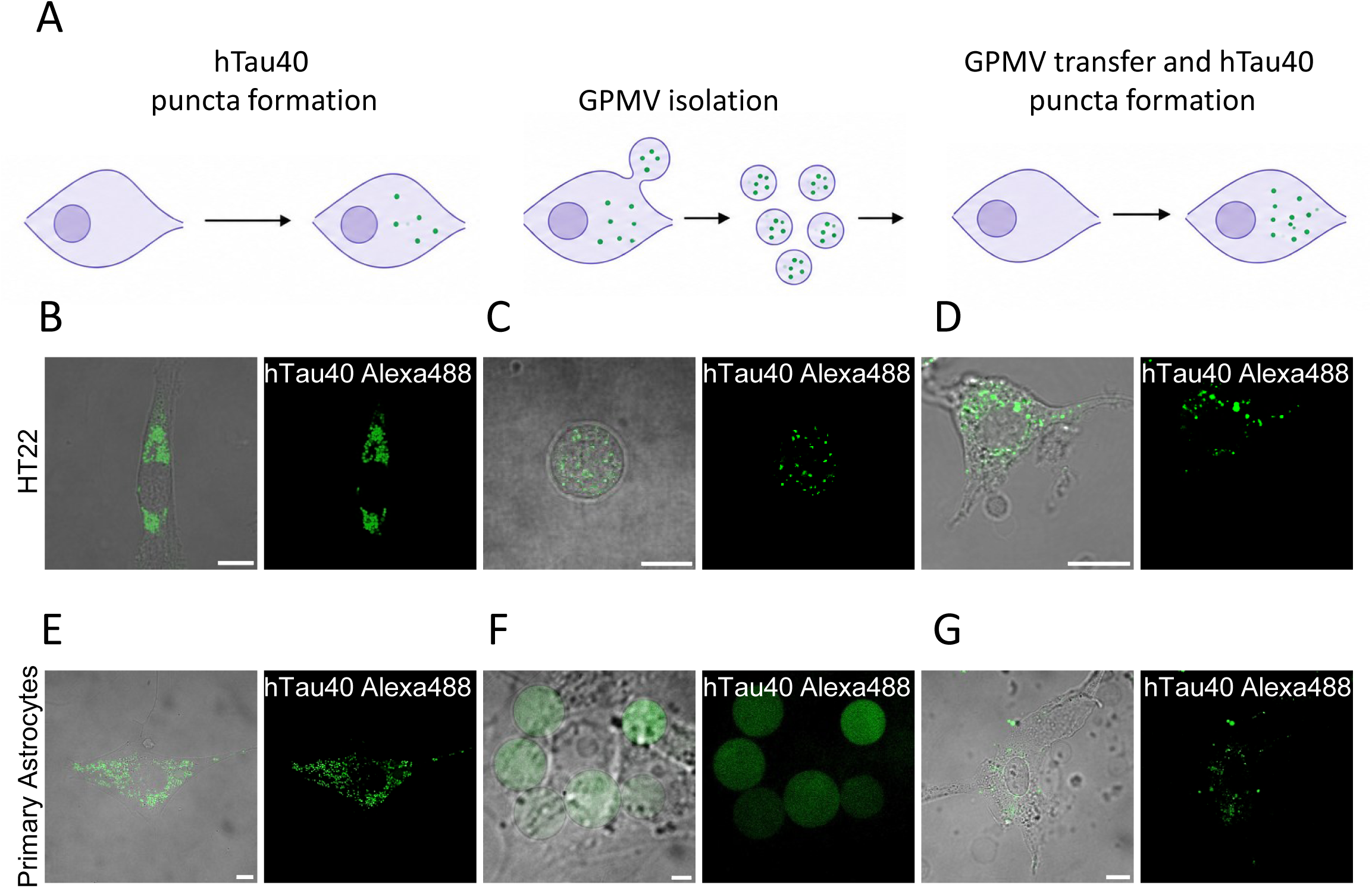
GPMV-mediated transfer of hTau40 induces intracellular puncta formation in recipient HT22 cells and primary cortical astrocytes. **(A)** Schematic illustration of the experimental workflow. Donor cells were incubated with hTau40-Alexa488 for 24 h to induce intracellular puncta formation, followed by induction and isolation of giant plasma membrane vesicles (GPMVs) over 12 h. Isolated GPMVs were subsequently transferred to untreated recipient cells, which were incubated for a further 24 h before imaging to assess the formation of intracellular puncta. **(B–D)** Representative merged brigshtfield/fluorescence images with the corresponding hTau40-Alexa488 fluorescence channel, illustrating the workflow in HT22 cells. **(B)** Donor HT22 cells containing intracellular hTau40-Alexa488 puncta. **(C)** GPMVs isolated from donor HT22 cells. **(D)** Recipient HT22 cells following GPMV transfer, showing intracellular hTau40 puncta. **(E–G)** Representative merged brightfield/fluorescence images with the corresponding hTau40-Alexa488 fluorescence channel, illustrating the same workflow in primary cortical astrocytes. **(E)** Donor astrocytes containing intracellular hTau40-Alexa488 puncta. **(F)** GPMVs isolated from donor astrocytes. **(G)** Recipient astrocytes following GPMV transfer, showing intracellular hTau40 puncta. Comparable observations in both cell types indicate that GPMV-mediated hTau40 transfer and puncta formation are not restricted to a single cell type. Scale bar: 10 µm.

We next asked whether this phenomenon is limited only to immortal cell lines such as HT22. Primary cortical astrocytes exposed to hTau40 similarly developed intracellular tau puncta, and GPMVs generated from these donor astrocytes retained hTau40 fluorescence (Fig. 8E, F). Following application to untreated recipient astrocytes, intracellular fluorescent tau puncta were again detectable after 24 h (Fig. 8G). The reproducibility of this transfer in primary astrocytes indicates that GPMV-mediated hTau40 transfer is not restricted to the immortalised HT22 cell model. Together, these experiments establish a reductionist plasma-membrane-vesicle system in which pre-existing intracellular hTau40 assemblies can be captured during vesiculation and subsequently transferred to recipient neural cells. Because GPMVs are experimentally induced membrane blebs rather than physiological extracellular vesicles, these observations do not establish an endogenous EV pathway. Instead, they demonstrate that low-mobility full-length hTau40 assemblies remain compatible with membrane enclosure and intercellular delivery.

## Discussion

Membrane lipids are increasingly recognised as modulators of tau assembly and toxicity, yet much of the mechanistic understanding of tau-membrane interactions derives from truncated repeat-domain constructs rather than full-length tau[28–30]. Here, neuronal lipid composition reshapes hTau40 aggregation and fibril mechanics, while tau assembly state governs membrane binding and remodelling. Notably, oligomers showed strong membrane recruitment, whereas greater lipid reorganisation accompanied actively assembling tau, indicating that membrane affinity and remodelling capacity are distinct properties. The lipid-dependent behaviour of hTau40 argues against the model that lipids can either promote or inhibit fibrillation. While anionic membranes can facilitate tau assembly[14, 30, 31], membrane interactions can also redirect full-length tau into stable tau-phospholipid complexes rather than canonical fibrils[17]. The mismatch between the ThT signal and endpoint morphology suggests that lipid composition redistributes hTau40 among distinct assembly states rather than simply altering fibril mass (Fig. 1). This is particularly relevant to PS, whose effects on full-length tau depend strongly on acyl-chain composition - saturated PS can promote aggregation, whereas unsaturated PS species suppress 1N4R and 2N4R fibrillation under heparin-free conditions[32]. Together with the polymorphic assemblies reported on anionic membranes[29], these findings support lipid specificity as a regulator of the tau assembly landscape. Indeed, neuronal membrane composition is known to undergo significant changes over time, thereby shaping the local lipid stoichiometry, which can influence the assembly state of tau[33, 34].

Lipid-dependent assembly is also reflected in fibril bending mechanics. Persistence length indicates resistance to bending and is sensitive to intermolecular packing and supramolecular architecture [21]. The increased flexibility of several lipid-associated fibrils therefore indicates altered mesoscale organisation, although its structural basis cannot be resolved without higher-resolution analysis. Broad bending distributions further reveal heterogeneity obscured by population-averaged persistence lengths[20], consistent with the structural polymorphism of heparin-induced full-length 2N4R tau[35]. Even membrane-free hTau40 segregated into mechanically distinct populations, supporting the view that tau fibrils form heterogeneous polymorphic ensembles rather than a single structural species[36]. Thus, membrane-dependent tau polymorphism results in a distinct mechanical phenotype (Fig. 2, 3). Indeed, distinct tau conformers can propagate stably in cells and mice and underlie different tauopathies[7, 37]. Membrane recruitment of tau showed distinct binding only on TBE, and not BPC:BPS (Fig. 4), despite similar surface potentials, consistent with models in which local anionic-lipid density, lateral organisation, and membrane-dependent tau conformation govern productive engagement[28, 38, 39]. Similar observations have also been shown to drive binding in the case of Aβ and αSynuclein[40–42]. At matched concentration, preformed oligomers accumulated more rapidly than the evolving hTau40 pool (Fig. 5), supporting oligomerisation as a gain in membrane avidity[16]. However, strong binding did not directly predict membrane-order perturbation: ongoing hTau40 assembly produced pronounced lipid reorganisation, whereas oligomers remained membrane-active at much lower concentration (Fig. 6B). Together, these findings suggest that membrane affinity and remodelling capacity are two distinct properties of tau, with oligomerisation favouring membrane recruitment and continued higher-order assembly, potentially amplifying membrane disruption. Such membrane perturbation may be particularly relevant to neuronal proteostasis, given evidence that endolysosomal membrane perforation can facilitate lysosomal access to internalised cytosolic α-synuclein fibrils and promote their pathological aggregation and associated neuronal toxicity [43]. We next asked whether an analogous relationship is apparent after tau encounters intracellular membranes. Following internalisation, hTau40 formed low-mobility puncta that strongly associated with lysosomes. Although polyanionic cofactors and intracellular crowding can promote the maturation of tau condensates toward arrested or aggregate-like states[44], negligible FRAP recovery alone does not establish a solid phase. Lysosomal association is nevertheless relevant because internalised tau can accumulate within endolysosomal compartments and perturb lysosomal integrity[45, 46]. Here, Lyso-Flipper and JIND-Mor revealed complementary changes in lysosomal membrane packing/mechanical state and microviscosity[25, 26], extending the reciprocal tau-membrane interaction observed in vitro to an intracellular membrane compartment (Fig. 7).

As a separate extension, we asked whether persistent intracellular hTau40 assemblies remain compatible with membrane enclosure and intercellular transfer. GPMVs represent a reductionist plasma membrane-derived vesicle system rather than a physiological EV model[47]. The transfer observed in both HT22 cells and primary astrocytes establishes that low-mobility hTau40 assemblies can remain compatible with membrane-mediated delivery (Fig. 8). Still, it does not demonstrate selective cargo sorting, endogenous EV biogenesis, or templated conversion of recipient tau. This finding parallels the growing evidence that brain- and cell-derived extracellular vesicles carry seed-competent, structurally defined tau filaments capable of templating aggregation in recipient cells and propagating tau pathology between neurons[48–52].

Taken together, these findings support a reciprocal relationship in which membrane composition selects tau assembly states, while tau assembly dynamics determine the extent of membrane remodelling. Notably, strong membrane recruitment does not necessarily predict bilayer reorganisation. This principle may extend to the endolysosomal interface, although the underlying molecular connection remains unresolved. Future studies should determine whether lipid-dependent tau states differ in fragmentation, seeding, or endolysosomal escape.

### Experimental procedures Materials

Total brain extract (TBE), brain phosphatidylcholine (BPC), brain phosphatidylserine (BPS), PEG-biotin, and NBD 460-PE were obtained from Avanti Polar Lipids. Atto 647N-PE and Thioflavin T (ThT) were obtained from Sigma-Aldrich. Alexa Fluor 488 and Alexa Fluor 647 protein labelling kits were obtained from Invitrogen (Thermo Fisher Scientific). High-molecular-weight heparin was obtained from MP Biomedicals. Trimethylammonium-diphenylhexatriene (TMA-DPH) was obtained from Thermo Fisher Scientific. LysoTracker was obtained from Invitrogen, and LysoFlipper^TR^ was obtained from Spirochrome. Avidin and chloroform were obtained from Sigma-Aldrich. DMEM GlutaMAX, fetal bovine serum (FBS), antibiotic-antimycotic solution, trypsin-EDTA, and Hanks’ Balanced Salt Solution (HBSS) were obtained from Gibco, Thermo Fisher Scientific. DNase I was obtained from Sigma-Aldrich. Protease inhibitor tablets were obtained from Roche. UranyLess staining solution was obtained from Electron Microscopy Sciences (EMS). Carbon type-B, 300-mesh copper transmission electron microscopy grids were obtained from Ted Pella Inc. Chambered coverslips and imaging dishes were obtained from Ibidi.

### Ethical statement

All animal experiments were conducted in accordance with the guidelines of the Committee for the Purpose of Control and Supervision of Experiments on Animals (CPCSEA) (Registration No. 1634/GO/ReRcBiBt/S/12/CPCSEA; Date of Registration: 16.05.2023). The experimental protocols were approved by the Institutional Animal Ethics Committee (IAEC) of the National Institute of Science Education and Research (NISER), Bhubaneswar, India, and all procedures were performed in compliance with institutional and national ethical guidelines (Ethical Approval No. NISER/SBS/AH-322).

### Protein expression, purification, and fluorescent labelling

The plasmid encoding full-length human tau (hTau40) was obtained from Addgene (tau/pET29b, a gift from Peter Klein; Addgene plasmid #16316; RRID: Addgene_16316). The plasmid was propagated in *Escherichia coli* DH5α, purified, and subsequently transformed into *E. coli* BL21(DE3) cells for protein expression. Cultures were grown at 37°C to mid-logarithmic phase, and protein expression was induced by the addition of 1 mM isopropyl β-D-1-thiogalactopyranoside (IPTG; GoldBio) for 3-4 h. Cells were harvested by centrifugation and processed for protein purification as previously described [53], with the modifications outlined below. Cell pellets were resuspended in ice-cold lysis buffer containing 20 mM MES-KOH (pH 6.5), 1 mM EGTA, 0.2 mM MgCl₂, 5 mM dithiothreitol (DTT), 1 mM phenylmethylsulfonyl fluoride (PMSF), and EDTA-free protease inhibitor tablets (Roche). Cells were lysed by sonication using a Qsonica Q700 sonicator equipped with a 13 mm probe tip at 50% amplitude with 5 s on/10 s off pulses for approximately 15-20 min while maintained on ice. Following lysis, NaCl was added to a final concentration of 500 mM, and the lysate was heated at 100°C for 20 min. The heat-treated lysate was subjected to ultracentrifugation at 127000 × *g* for 40 min at 4°C to remove precipitated proteins and cellular debris. The resulting supernatant was dialysed overnight against cation-exchange equilibration buffer containing 20 mM MES, 50 mM NaCl, 1 mM EGTA, 1 mM MgCl₂, 2 mM DTT, and 0.1 mM PMSF (pH 6.5) using SnakeSkin™ dialysis tubing. The dialysate was subsequently clarified by ultracentrifugation to remove insoluble material before chromatographic purification. The clarified sample was loaded onto a HiTrap SP HP cation-exchange column (Cytiva) equilibrated with the corresponding equilibration buffer using an ÄKTA pure FPLC system (Cytiva). Bound protein was eluted using a linear gradient of elution buffer containing 20 mM MES, 1 M NaCl, 1 mM EGTA, 1 mM MgCl₂, 2 mM DTT, and 0.1 mM PMSF (pH 6.5). Fractions containing hTau40 were identified by UV absorbance and confirmed by SDS-PAGE, pooled, and concentrated using a Vivaspin™ Turbo 15 centrifugal concentrator with a 10 kDa molecular-weight cutoff (Sartorius). The concentrated protein was subsequently dialysed against gel-filtration buffer using the same dialysis tubing and subjected to size-exclusion chromatography on a HiPrep 16/60 Sephacryl S-200 HR column (Cytiva) connected to the ÄKTA pure system. The column was equilibrated with gel-filtration buffer consisting of PBS containing 1 mM DTT (pH 7.4). Fractions corresponding to the hTau40 peak were identified by UV absorbance and SDS-PAGE, pooled, and the protein concentration was determined using the bicinchoninic acid (BCA) assay. Purified hTau40 was aliquoted and stored at -80°C until further use. Purified hTau40 was fluorescently labelled with Alexa Fluor 488 or Alexa Fluor 647 using the corresponding Protein Labelling Kits (Invitrogen, Thermo Fisher Scientific; A10235 and A20173, respectively) according to the manufacturer’s instructions. Unreacted fluorescent dye was removed using the purification resin supplied with the respective labelling kit. Labelled hTau40 was subsequently stored under the same conditions as the unlabelled protein until use. Unless otherwise specified, fluorescently labelled hTau40 was mixed with unlabelled hTau40 at a 1:10 labelled-to-unlabelled protein ratio for subsequent experiments.

### Preparation of large unilamellar vesicles(LUVs)

LUVs were generated following a previously reported procedure[41], with modifications. Briefly, 1 mM lipid stocks were first evaporated under a gentle stream of nitrogen to form a thin lipid film, then vacuum-dried for 1 h. The dried lipids were resuspended in 1 mL of PBS (pH 6.5) and allowed to hydrate for ∼15 min in a water bath maintained above the phase-transition temperature of the corresponding lipid composition. The hydrated suspension was vortexed for 4-5 min to facilitate the formation of multilamellar vesicles. These vesicles were subsequently sonicated using a Sonics VCX130 equipped with a 3 mm probe, operated at 50% amplitude with alternating 10s on and 5s off pulses for a total sonication time of 3 min. The resulting LUV preparations were characterised for hydrodynamic diameter and zeta potential using a Malvern Zetasizer Nano ZS90. The mean vesicle diameter was ∼150 nm.

### ThT aggregation kinetics and analysis

Tau aggregation kinetics were monitored using a thioflavin T (ThT) fluorescence assay adapted from [54] with modifications. The reaction buffer consisted of PBS (pH 6.5) supplemented with 1 mM DTT and 0.05% (w/v) sodium azide. Purified hTau40 was thawed, allowed to equilibrate to room temperature, and briefly centrifuged to remove air bubbles. Protein concentration was determined using the BCA assay. ThT was added at a 1:1 molar ratio relative to hTau40, and aggregation was initiated by the addition of high-molecular-weight heparin (∼17 kDa; MP Biomedicals) at a hTau40:heparin molar ratio of 4:1. For experiments performed in the presence of membranes, LUVs were included at a hTau40:lipid molar ratio of 1:4. Reaction mixtures were dispensed into black, flat-bottom 96-well microplates (Greiner Bio-One), and ThT fluorescence was monitored using a BMG CLARIOstar microplate reader in bottom-read mode. Fluorescence was measured at an excitation wavelength of 440 nm and an emission wavelength of 485 nm. Measurements were performed at 37°C with orbital shaking at 200 rpm, with fluorescence readings collected at 15 min intervals for a total duration of 125 h. The resulting fluorescence traces were fitted to a four-parameter Boltzmann sigmoidal model:

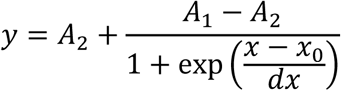

where *A*_1_ and *A*_2_ represent the lower and upper asymptotes of the fitted curve, respectively, *x*_0_ denotes the inflexion point, and (dx) is the slope factor describing the steepness of the transition. The maximum aggregation growth rate was calculated from the maximum slope of the fitted Boltzmann function:

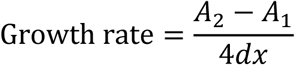

The upper asymptote *A*_2_ was taken as the saturation value of the aggregation reaction. Thus, *A*_2_ represents the fitted plateau fluorescence intensity for each experimental condition. The fitted kinetic parameters, including the growth rate and saturation value, were used to quantitatively compare aggregation behaviour across different membrane conditions.

### Electron microscopy sample preparation and imaging

Transmission electron microscopy (TEM) samples were prepared following previously reported procedures[17, 41], with modifications. Briefly, 20 µL of the sample was deposited onto carbon type-B, 300-mesh copper grids (Ted Pella Inc.) and allowed to adsorb for 30 min. The grids were then blot-dried and rinsed with ultrapure water before negative staining with UranyLess staining solution (Electron Microscopy Sciences). A drop of the staining solution was applied to each grid, which was then incubated for 2 min, after which the grids were briefly rinsed with ultrapure water and blot-dried. TEM imaging was performed using a JEOL JEM F200 transmission electron microscope.

### Persistence length and mechanical property analysis

The mechanical properties of tau fibrils were assessed from contour length (*L*) and end-to-end distance (*R*) measurements obtained by manually tracing fibrils in TEM micrographs using Simple Neurite Tracer (SNT) in Fiji/ImageJ. For each fibril, a dimensionless bending parameter, η, was calculated as:

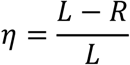

Values approaching zero correspond to relatively straight fibrils, whereas increasing η values indicate greater bending and conformational flexibility[20]. For each experimental condition, the mean-square end-to-end distance (R²) was plotted as a function of contour length (*L*) and fitted using the two-dimensional worm-like chain (2D WLC) model[20, 55]:

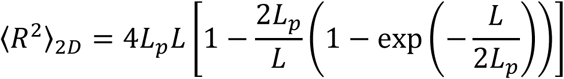

where *L*_*p*_is the persistence length. The model was fitted by nonlinear least-squares regression. The uncertainty in *L*_*p*_was estimated by bootstrap resampling (1000 iterations); the reported ± error represents half the width of the 95% confidence interval derived from the bootstrap distribution. To account for differences in fibril length, η distributions were represented as mass-weighted histograms (Fig. 3A-E). For each η bin, the histogram value was calculated as the sum of the contour lengths (Σ*L*, nm) of all fibrils falling within that bin, thereby weighting each fibril according to its relative mass contribution[20]. Histogram bin widths were determined independently for each condition using the Freedman-Diaconis[56]:

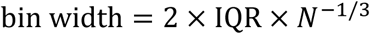

where IQR is the interquartile range of the η values, and N is the number of fibrils, giving bin counts of 8, 11, 9, 10, and 9 for hTau40, TBE+hTau40, BPC+hTau40, BPS+hTau40, and BPC-BPS+hTau40, respectively. The theoretical η distribution expected for a homogeneous fibril population was generated by Monte Carlo simulation of the 2D WLC model following [22]. For each experimentally measured fibril, 100,000 WLC conformations were generated using a segment length of 5 nm. At each segment junction, the bending angle was sampled from a Gaussian distribution with zero mean and a standard deviation given by:

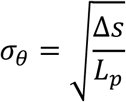

where *Δs* is the segment length. The resulting conformations were used to calculate the end-to-end distance and η for each simulated fibril. The simulated distributions were mass-weighted according to contour length and area-normalised to the corresponding experimental histograms for visual comparison. The theoretical distributions were generated using the independently fitted *L*_*p*_without adjusting additional parameters to reproduce the experimental histogram shape. For the hTau40 condition, the presence of mechanically distinct fibril subpopulations was evaluated using Gaussian Mixture Models (GMMs) applied to individual fibril η values (*N* = 236). Models containing one to six Gaussian components were evaluated, with 20 independent initialisations for each model. Model selection was performed using the Bayesian Information Criterion (BIC)[23], with the model exhibiting the lowest BIC selected as the preferred model. The two-component model yielded the global minimum BIC, with ΔBIC = 37.2 relative to the one-component model, whereas models containing three or more components were progressively disfavoured. The separation threshold between the two fitted populations was defined as the crossover point of their mixing-proportion-weighted Gaussian density functions, yielding η* = 0.369. Fibrils with η < η* were assigned to Population 1 (*N* = 121), whereas fibrils with η ≥ η* were assigned to Population 2 (*N* = 115). The two subpopulations were subsequently analysed independently by fitting the 2D WLC model to their respective ⟨R²⟩ versus *L* distributions (Fig. 3F). Persistence lengths and associated uncertainties were determined using the same nonlinear least-squares and bootstrap procedures described above. The bending modulus, *C*_*B*_, was calculated from the persistence length according to the relationship described by [20, 55]:

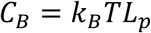

where *k*_*B*_ is the Boltzmann constant (1.380649 × 10⁻²³ J K⁻¹), *T* is the absolute temperature (310 K; 37°C), and *L*_*p*_was converted from nanometres to meters before calculation. The resulting bending modulus was expressed in N·m², with uncertainty propagated directly from the uncertainty in *L*_*p*_. All numerical analyses, nonlinear fitting, statistical modelling, and Monte Carlo simulations were performed in Python using NumPy, SciPy, scikit-learn, openpyxl, and Matplotlib. A fixed random seed was used for stochastic analyses to ensure reproducibility.

### Preparation of fluorescently labelled GUVs and confocal imaging

Giant unilamellar vesicles (GUVs) were prepared using a gel-assisted method adapted from [41, 57] with modifications. Briefly, a 5% (w/w) polyvinyl alcohol (PVA) solution was prepared in deionised water, and 300 µL of the solution was evenly spread onto a plastic Petri dish and dried at 50°C for 30 min. The dried PVA-coated Petri dishes were then exposed to UV light for 15 min to promote dewetting of the subsequently deposited lipid film. A lipid solution in chloroform (5 mM), containing 0.1 mol% Atto647-PE or NBD460-PE and 0.1 mol% PEG-biotin, was subsequently spread onto the PVA-coated surface (20-25 µL). The lipid-coated Petri dishes were placed under vacuum for 45-60 min to remove residual solvent. The lipid films were then hydrated with PBS (pH 6.5) for 15 min, and the resulting GUVs were gently detached from the PVA layer and transferred to microcentrifuge tubes using a pipette. For imaging experiments, ibidi chambered coverslips were first incubated with avidin (0.1 mg/mL) for 10 min, then washed with PBS (pH 6.5). The GUV suspension was then introduced into the avidin-coated chamber to facilitate GUV immobilisation through the biotin-avidin interaction. GUVs were incubated with 10 µM hTau40, containing both fluorescently labelled and unlabelled forms, at a 1:10 molar ratio. Sodium azide was included at a final concentration of 0.05% (w/v). Samples were monitored by confocal microscopy (Olympus FLUOVIEW FV3000) for up to 3 days to assess hTau40 association with the GUV membranes and associated changes in membrane organisation.

### Preparation and characterisation of tau oligomers and aggregates

Tau oligomers and aggregates were prepared as previously described by [58–61]. Briefly, Purified hTau40 was incubated with polyanionic heparin (∼17.5 kDa) at a hTau40:heparin molar ratio of 4:1 in 1X PBS containing 2 mM DTT, 0.01% sodium azide and 1X protease inhibitor cocktail for 12 h at 37°C. Following incubation, glutaraldehyde was added to a final concentration of 0.01%, and the reaction mixture was incubated for an additional 10 min to stabilise the oligomeric species. Stabilised tau oligomers were buffer-exchanged into 1X PBS using 5-kDa centricons by centrifugation at 3,800 rpm at 4 °C. For aggregate preparation, hTau40 was incubated with heparin at a 4:1 molar ratio in the presence of 1 mM DTT at 37°C for 72 h to promote fibril formation. The resulting tau preparations were characterised by SDS-PAGE and Thioflavin S (ThS) fluorescence (Fig. S4, A-B). For ThS analysis, tau species were mixed with ThS (Sigma Aldrich) prepared in ammonium acetate buffer at a hTau40:ThS ratio of 1:4. Samples were prepared in triplicate along with buffer blanks, incubated for 10 min in the dark at room temperature, and fluorescence was measured using a Tecan Spark microplate reader at 440 nm excitation and 521 nm emission. For ultrastructural characterisation, tau oligomers and aggregates were diluted to 2 µM, and 10 µL of each sample was applied to 400-mesh carbon-coated copper grids (Ted Pella) for 90 s. Grids were rinsed with filtered Milli-Q water for 30 s, stained with 2% uranyl acetate for 2 min, and air-dried before overnight incubation at 37°C. Samples were subsequently examined by transmission electron microscopy (JEM-1400Plus) to assess the morphology of the different tau assembly states (Fig. S4, C-D). Following confirmation of the respective assemblies, oligomeric and aggregated tau preparations were concentrated and stored at −80°C until further use.

### Fluorescence anisotropy assay

Fluorescence anisotropy measurements were performed using LUVs prepared as described above. Trimethylammonium-diphenylhexatriene (TMA-DPH) was dissolved in DMSO to prepare a 2 mM stock solution. LUVs were incubated with TMA-DPH at a TMA-DPH:lipid molar ratio of 1:100 for 30 min in the dark. Following probe incorporation, hTau40 was added at a final concentration of 50 µM together with high-molecular-weight heparin at a hTau40:heparin molar ratio of 4:1. hTau40 oligomers were analysed separately at a final concentration of 1 µM. The resulting mixtures were transferred to black, flat-bottom microplates (Brand), and fluorescence anisotropy was measured using a BMG CLARIOstar microplate reader in bottom-read mode. Measurements were performed at 37°C for 125 h, with readings collected at 1 h intervals using excitation and emission wavelengths of 360 and 430 nm, respectively. Fluorescence anisotropy was calculated according to:

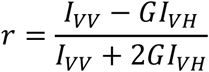

where *I*_*VV*_ and *I_VH_* represent the fluorescence intensities measured with vertically polarised excitation and vertically or horizontally polarised emission, respectively. *G*, defined as *G* = 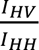 is the grating correction factor used to account for wavelength-dependent differences in the instrument’s polarisation response. The resulting anisotropy values were plotted as a function of time and fitted by linear regression:

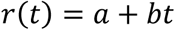

where *a* is the intercept and *b* is the slope, representing the rate of change in membrane anisotropy with time. Positive and negative values of *b* indicate increasing and decreasing anisotropy, respectively.

### Mammalian cell culture, tau treatment, and lysosomal colocalization

Mouse hippocampal neuronal HT22 cells were cultured in DMEM GlutaMAX supplemented with 10% fetal bovine serum (FBS) and 1% antibiotic-antimycotic solution at 37°C in a humidified incubator containing 5% CO₂. Early-passage cells (passages 1-10) were used for all experiments. For Tau treatment, the culture medium was replaced with fresh medium containing 1 µM monomeric hTau40 and high-molecular-weight heparin at a hTau40:heparin molar ratio of 4:1. Cells were incubated with Tau for 24 h. For lysosomal colocalization analysis, cells were subsequently incubated with 70 nM LysoTracker (Invitrogen) for 30 min. Cells were then imaged using an Olympus FLUOVIEW FV3000 confocal microscope. For live-cell imaging, cells were maintained under controlled environmental conditions using a TOKAI HIT stage-top incubator during image acquisition.

### Fluorescence recovery after photobleaching (FRAP)

FRAP measurements were performed on hTau40-treated HT22 cells using an Olympus FLUOVIEW FV3000 confocal microscope. Before photobleaching, images were acquired at attenuated laser intensity to establish the pre-bleach fluorescence baseline. hTau40-Alexa647 was photobleached using an OBIS 640 LX laser at 100% laser power for 10 s. Following photobleaching, the laser intensity was returned to the attenuated acquisition setting, and fluorescence recovery was monitored for 600 s using time-lapse imaging. Five independent recovery measurements were obtained for each experimental condition and averaged to generate the mean fluorescence recovery curve. The recovery curves were fitted using a single-exponential recovery model:

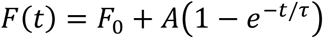

where *F*(*t*) is the fluorescence intensity at time *t*, *F*_0_ is the fluorescence intensity immediately after photobleaching, *A* is the recovery amplitude, and *τ* is the characteristic recovery time constant. The asymptotic fluorescence intensity at complete recovery was calculated as: *F*_∞_ = *F*_0_ + *A*. The fitted recovery curves were used to characterise the extent and kinetics of fluorescence recovery following photobleaching.

### Fluorescence lifetime imaging microscopy (FLIM)

FLIM measurements were carried out on a MicroTime 200 (MT200) time-resolved confocal microscope (PicoQuant, Berlin, Germany) coupled to a time-correlated single-photon counting (TCSPC) module. Samples were excited using pulsed laser illumination at either 488 or 519 nm, depending on the fluorophore being examined. The excitation power at the sample was maintained between 10 and 30 nW, as measured following the dichroic mirror. Approximately 2 × 10⁴ HT22 cells were plated on glass-bottom dishes and imaged using a 60X water-immersion objective. The appropriate dichroic mirror was selected according to the excitation wavelength. A 50 µm pinhole was used to suppress out-of-focus emission, while the fluorescence was split 50:50 with a beam splitter and detected by single-photon avalanche diodes (SPADs). Following 24 h of treatment with hTau40 under the conditions described above, cells were incubated with 1 µM Lyso-Flipper-TR (Spirochrome) for 30 min to measure lysosomal membrane order. For lysosomal viscosity measurements, cells were instead incubated with 1 µM JIND-Mor for 30 min under the same conditions. Fluorescence decay data were collected using the TCSPC module and analysed with FLIMfit software developed at Imperial College London. Fluorescence lifetime parameters obtained from the fitted decay profiles were used for subsequent quantitative analysis. All measurements were acquired using identical imaging and acquisition settings across experimental conditions. Experiments were performed at room temperature using a minimum of three independent biological replicates.

### Giant plasma membrane vesicles (GPMV) isolation

GPMVs were generated from HT22 cells using a chemical vesiculation procedure[27]. Cells were seeded at 2 × 10^5^ cells per well in 6-well plates and cultured until they reached ∼60-80% confluency. Cells were maintained under identical culture conditions and incubated overnight before vesiculation. To generate hTau40-containing GPMVs, cells were treated with hTau40 under the conditions described above for 24 h before vesicle isolation. Before vesiculation, cells were washed twice with 1 mL of GPMV buffer containing 10 mM HEPES, 150 mM NaCl, and 2 mM CaCl₂ to remove residual medium containing any extracellular protein, non-adherent cells, and cellular debris. GPMV formation was induced by incubating the cells in GPMV buffer supplemented with 2 mM DTT and 0.07% (w/v) paraformaldehyde for 12-16 h at 37°C. Vesicle formation was monitored by light microscopy, with GPMVs identified as spherical, membrane-enclosed structures. Following vesiculation, the culture plates were gently tapped to detach surface-associated GPMVs, and the resulting supernatant was carefully collected. Intact cells and larger cellular debris were removed by centrifugation at 500 × g for 5 min, while GPMVs remained in the supernatant. The resulting GPMV suspension was collected and transferred to freshly cultured recipient HT22 cells at ∼50-60% confluency. Recipient cells were incubated with isolated GPMVs at 37°C for 24 h to allow uptake of the vesicles. Cells were subsequently processed for confocal imaging.

### Primary cortical astrocyte culture and GPMV transfer

Primary cortical astrocytes were isolated from postnatal day 0-1 (P0-P1) Sprague-Dawley rat pups. Following brain removal, tissues were immediately transferred to ice-cold calcium- and magnesium-free Hanks’ Balanced Salt Solution (HBSS, Gibco) containing 10 mM glucose and HEPES (Sigma-Aldrich). The cortical regions were dissected and enzymatically dissociated with 0.25% trypsin-EDTA (Gibco) and 150 U/mL DNase I (Sigma-Aldrich) for 15 min at 37°C. Enzymatic digestion was terminated by adding 10% fetal bovine serum (FBS; Gibco). The tissue was then mechanically dissociated by gentle trituration and centrifuged at 1000 rpm for 5 min at 4°C. The resulting cell pellet was resuspended in DMEM (Sigma-Aldrich) containing 10% FBS and 1% antibiotic-antimycotic solution (Gibco), and cells were seeded at cells per T25 flask. Cultures were maintained at 37°C in a humidified incubator with 5% CO₂. After ∼2 weeks, when the cultures reached confluency, the flasks were subjected to orbital shaking at 220 rpm for 18 h to remove loosely adherent microglia and oligodendrocyte precursor cells. Following shaking, the cultures were washed with 1 mL warm 1X PBS to remove remaining suspended cells and debris. The adherent astrocyte monolayer was then detached using trypsin and replated onto poly-D-lysine-coated imaging dishes (0.1 mg/mL) for live-cell imaging or onto 100-mm dishes for GPMV generation and collection[62, 63]. GPMVs were generated and isolated as described above, with the DTT concentration increased to 4 mM. For the generation of hTau40-containing astrocyte GPMVs, donor astrocytes were treated with hTau40 under the conditions described previously, 24 h before GPMV induction. The resulting hTau40-containing GPMVs were isolated using the same procedure described above and transferred to recipient astrocyte cultures maintained at ∼50-60% confluency. Recipient cultures were incubated with isolated GPMVs at 37°C for 24 h to allow vesicle uptake, then processed for confocal imaging.

### Image processing and statistical analysis

Image processing and quantitative image analysis were performed using Fiji/ImageJ unless otherwise specified. Graphs were generated using OriginPro. Statistical analyses were performed using the Kruskal-Wallis test followed by Dunn’s multiple-comparisons post hoc test, unless otherwise indicated in the corresponding analysis. Statistical significance was defined as *p* < 0.05.

## Data availability

All the data are available within the main article and the supporting information files.

## Acknowledgements

M.S. acknowledges the financial support received from the Science & Engineering Research Board (SERB) (Grant No. EMR/2017/004513), Department of Biotechnology, Govt. of India (Grant No. BT/PR/21226/ MED/122/41/2016) and DBT-Wellcome Trust India Alliance Intermediate Fellowship (Grant No. - IA/I/20/2/505212). S.G. acknowledges SERB (Grant no. SRG/2022/000117), Indian Council of Medical Research (ICMR) (Grant no. IIRP-2023-0585) and DBT (Grant no. BT/PR48748/MED/122/329/2023) for funding. M.S., A.S.M. and S.G. also acknowledge the Department of Atomic Energy (DAE) for the intramural financial support. A.K.M. thanks CSIR, Ministry of Science & Technology, Govt. of India, for the doctoral fellowship. We also thank Apurba Lal Koner for sharing JIND-Mor. We acknowledge Smita Desale for the help during hTau40 oligomer and aggregates preparation and characterisation.

