## Supplementary Data for "Neuronal lipid composition regulates distinct states of full-length tau assembly, mechanics and membrane interactions"

This supplementary information file contains supplementary figures S1–S6 and the corresponding legends for supplementary movies S1–S4

Figure S1

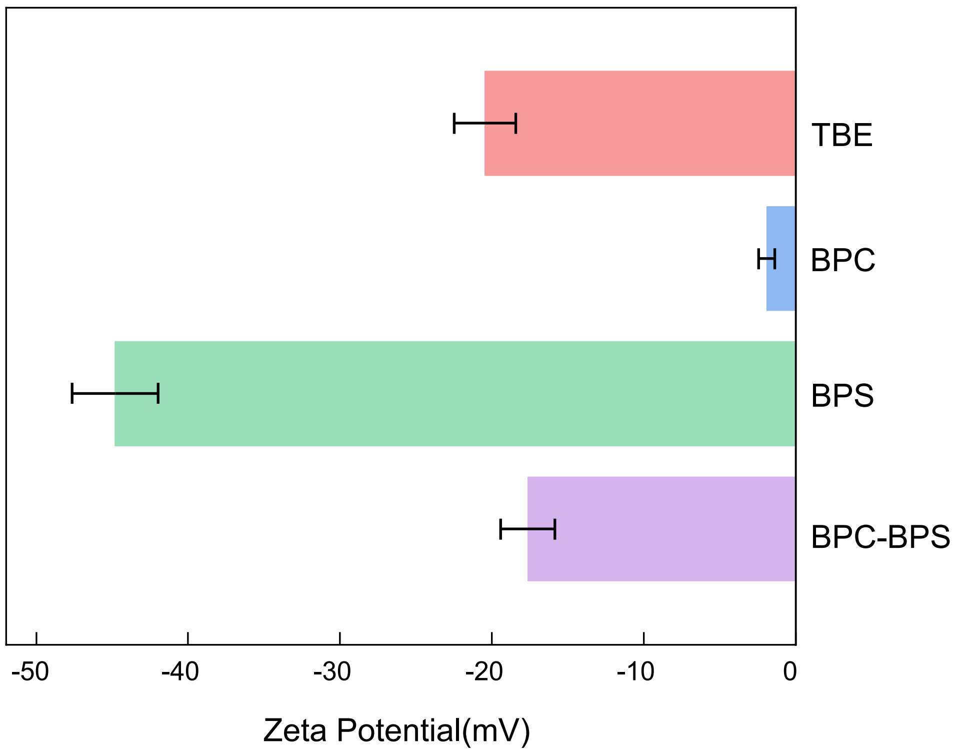

Supplementary Figure S1. Zeta potential of large unilamellar vesicles (LUVs) used across all membrane conditions

Zeta potential measurements of LUVs composed of TBE, BPC, BPS and BPC:BPS (6:4), used throughout all experiments. BPC vesicles exhibited a near-neutral zeta potential, whereas TBE, BPS, and BPC-BPS carried a net negative surface charge. Data represent mean ± SD from three independent experiments.

### Figure S2

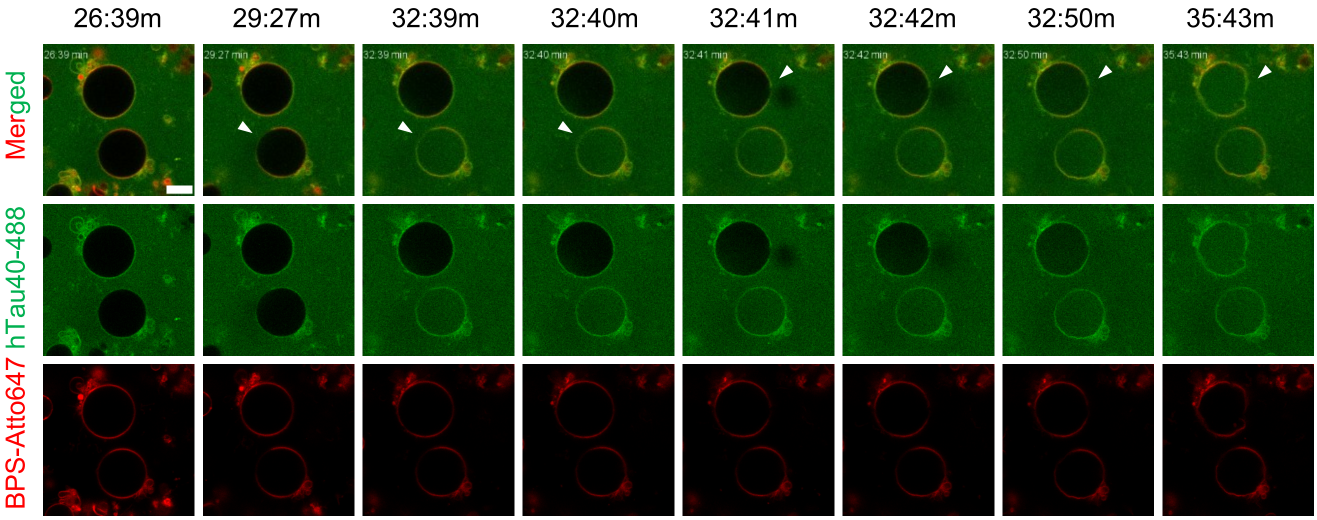

26:39m
29:27m
32:39m
32:40m
32:41m
32:42m
32:50m
35:43m
Merged
hTau40-488
BPS-Atto647

Supplementary Figure S2. hTau40 binds BPS giant unilamellar vesicles and induces membrane rupture within minutes

Time-lapse confocal images of hTau40-Alexa488 (10 µM) with heparin (tau:heparin, 4:1) incubated with BPS GUVs (Atto647), showing membrane binding followed by GUV rupture between approximately 26 and 36 min. The merged hTau40-Alexa488 and BPS-Atto647 channels are shown at the indicated time points. Scale bar: 10 µm.

Figure S3

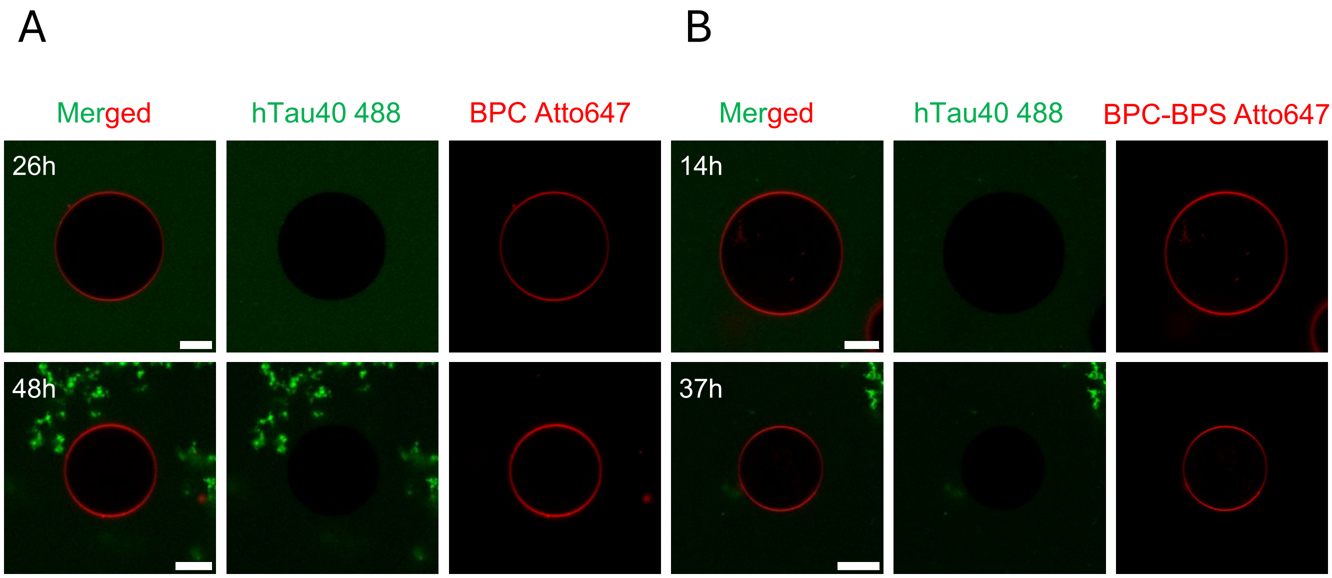
A
B
Merged
hTau40 488
BPC Atto647
Merged
hTau40 488
BPC-BPS Atto647
26h
48h
14h
37h

Supplementary Figure S3. hTau40 shows no detectable membrane binding to BPC or BPC-BPS giant unilamellar vesicles over extended incubation.

Representative confocal images of hTau40-Alexa488 (10 µM, with heparin, tau:heparin 4:1) incubated with **(A)** BPC GUVs (Atto647) at 26 h and 48 h, and **(B)** BPC-BPS GUVs (Atto647) at 14 h and 37 h. Merged, hTau40-Alexa488, and respective membrane channels are shown at the indicated time points. No detectable membrane binding of hTau40 to either GUV membrane was observed at any time point, consistent with the corresponding conditions shown in Figure 4:C-D. Scale bar: 10 µm.

### Figure S4

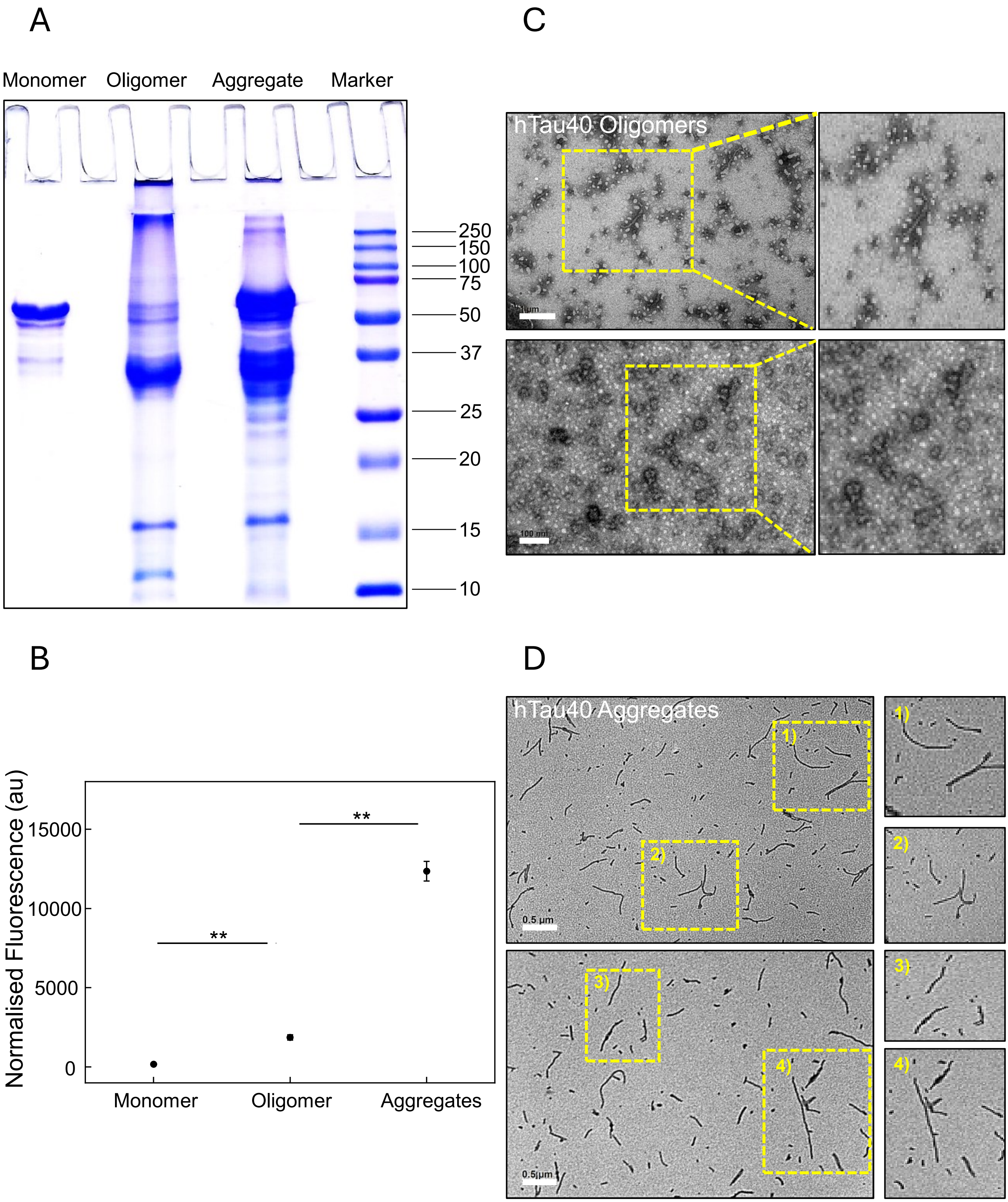
A
Monomer
Oligomer
Aggregate
Marker
250
150
100
75
50
37
25
20
15
10
B
Normalised Fluorescence (au)
15000
10000
5000
0
**
**
Monomer
Oligomer
Aggregates
C
hTau40 Oligomers
100 nm
D
hTau40 Aggregates
1)
2)
3)
4)
0.5 µm

Supplementary Figure S4. Biochemical and ultrastructural characterisation of distinct hTau40 assembly states

**(A)** Coomassie-stained 10% SDS-PAGE analysis of monomeric, oligomeric, and aggregated hTau40 preparations, showing distinct electrophoretic profiles of the different Tau assembly states. **(B)** Thioflavin S (ThS) fluorescence analysis of the corresponding hTau40 preparations, showing increased amyloid-associated fluorescence in oligomeric and aggregated samples relative to monomeric hTau40. Statistical significance was assessed using the Mann-Whitney U test; $p < 0.01$. **(C)** Transmission electron microscopy (TEM) images of negatively stained oligomeric hTau40, showing small, rounded particulate species. **(D)** TEM images of negatively stained aggregated hTau40, showing predominantly elongated fibrillar structures. Scale bars are indicated in the respective panels.

Figure S5

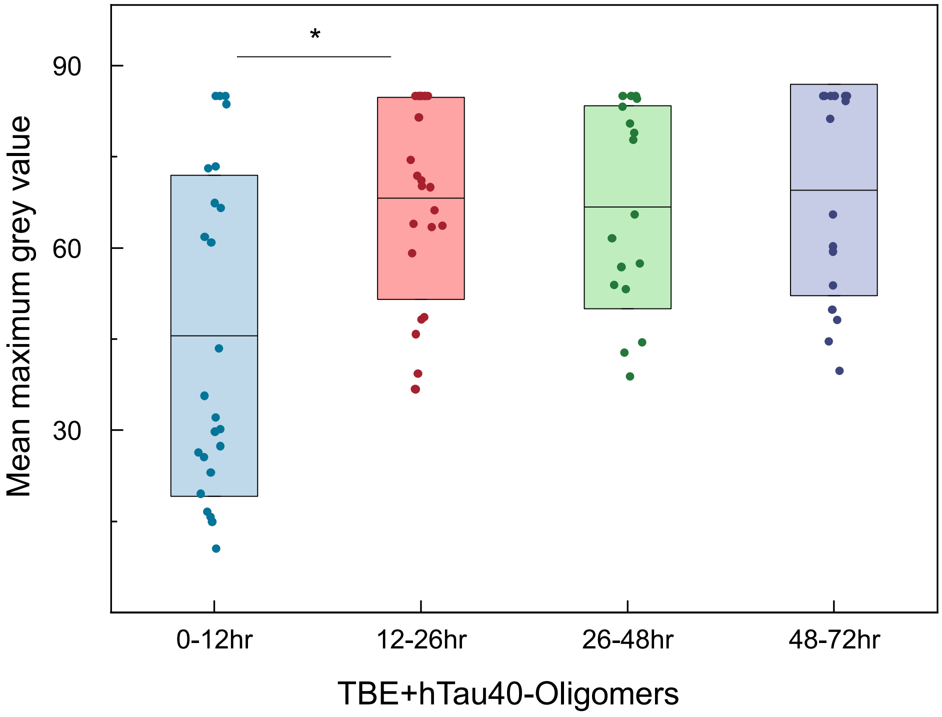

*
90
60
30
Mean maximum grey value
0-12hr
12-26hr
26-48hr
48-72hr
TBE+hTau40-Oligomers

Supplementary Figure S5. Maximum-fluorescence-intensity quantification confirms the progressive binding of hTau40 oligomers to TBE giant unilamellar vesicles.

Quantification of hTau40 oligomer binding to TBE GUVs, determined from the mean maximum grey value obtained by 360-point oval intensity profiling of the GUV membrane, binned across four time windows (0-12 h, 12-26 h, 26-48 h and 48-72 h). Approximately 20 GUVs were analysed per time bin. A significant increase in membrane-associated intensity was observed between 0-12 h and 12-26 h, consistent with the top 10% grey value quantification shown in Figure 6A. Data are presented as mean ± SD box plots with individual data points overlaid. Statistical significance was determined by the Kruskal-Wallis test with Dunn's post hoc analysis; * $p < 0.05$.

Figure S6

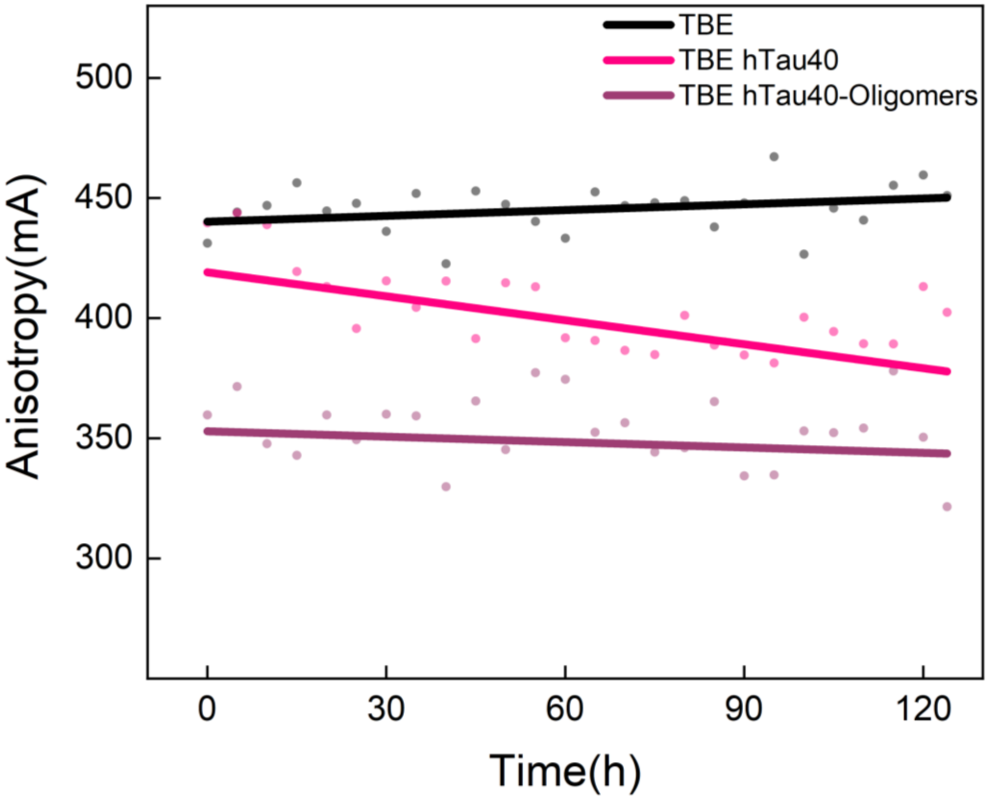
TBE
TBE hTau40
TBE hTau40-Oligomers
Anisotropy(mA)
500
450
400
350
300
0
30
60
90
120
Time(h)

Supplementary Figure S6. Raw anisotropy data over time for TBE membranes with and without hTau40.

Raw TMA-DPH steady-state anisotropy values over time for TBE alone, TBE with hTau40 (50µM with heparin, tau:heparin 4:1), and TBE with hTau40 oligomers (1µM without heparin), measured over 125 h. Solid lines represent linear fits to the raw data. The slopes of the fitted lines were used to calculate the anisotropy change rates presented in Figure 6B.

Supplementary Movie 1. Time-lapse recording of progressive hTau40 binding to a TBE giant unilamellar vesicle.

Time-lapse confocal recording of hTau40-Alexa488 (with heparin; tau:heparin, 4:1) binding to a TBE giant unilamellar vesicle (GUV; Atto647) from 6 to 22 h, demonstrating progressive membrane binding. Representative frames are shown in Figure 4A. Scale bar: 10µm

Supplementary Movie 2. Time-lapse recording of hTau40-induced rupture of a BPS giant unilamellar vesicle.

Time-lapse confocal recording of hTau40-Alexa488 (with heparin; tau:heparin, 4:1) binding to a BPS giant unilamellar vesicle (GUV; Atto647), demonstrating membrane binding followed by GUV rupture within the first hour of incubation. Representative frames are shown in Supplementary Figure S2. Scale bar: 10µm

Supplementary Movie 3. GPMV formation from an HT22 cell containing intracellular hTau40 puncta.

Time-lapse confocal recording of an HT22 cell containing intracellular hTau40-Alexa647 puncta during giant plasma membrane vesicle (GPMV) formation, demonstrating hTau40 puncta within both the donor cell and the attached GPMV. Scale bar: 10µm

Supplementary Movie 4. Isolated giant plasma membrane vesicle containing intracellular hTau40 puncta.

Time-lapse confocal recording of an isolated giant plasma membrane vesicle (GPMV) containing hTau40-Alexa488 puncta following isolation from a donor HT22 cell. Scale bar: 10µm
